# The role of the locus coeruleus in eye movements during perceptual decision making

**DOI:** 10.64898/2026.03.01.708911

**Authors:** Katerina Acar, Matthew A. Smith

**Affiliations:** Department of Biomedical Engineering, Carnegie Mellon University, Pittsburgh, PA, USA; Center for the Neural Basis of Cognition, Carnegie Mellon University and University of Pittsburgh, Pittsburgh, PA, USA; Neuroscience Institute, Carnegie Mellon University, Pittsburgh PA, USA

## Abstract

The locus coeruleus (LC) is the primary source of norepinephrine in the brain and has been implicated in the processes of attention, arousal, and perceptual decision making. Although prior work has linked transient LC activation to both sensory stimulus processing and motor processing, the precise contribution of LC to the distinct sensory and motor components of perceptual decisions remains unclear. Here, we recorded the spiking activity of single LC neurons in rhesus macaques while they performed a visual two-alternative forced-choice change detection task with a saccadic report, designed to cleanly dissociate sensory and motor contributions to LC activity. We found that the large majority of recorded neurons showed robust increases in response tightly locked to the choice saccade, while only a small fraction showed significant responses to the visual stimuli. Saccade-aligned LC responses did not vary with behavioral outcome, perceptual difficulty, reaction time, or session-wide fluctuations in perceptual sensitivity and criterion, indicating that LC motor-related signals were dissociated from perceptual performance. Together, these results demonstrated the existence of a subpopulation of LC neurons whose activity was tightly coupled to oculomotor output across both voluntary and involuntary eye movements during perceptual decision making, but were independent of perceptual decision accuracy. Our findings support a role for LC in facilitating motor preparation and execution in response to behaviorally significant sensory events.

## Introduction

Pioneering electrophysiological recordings from the locus coeruleus (LC) in awake animals reported that the presentation of a startling or otherwise salient sensory stimulus, or a noxious or preferred food presentation, evoked a transient burst of spikes now known as phasic activation (Aston-Jones and Bloom 1981; Foote et al. 1980; Grant et al. 1988). Subsequent work demonstrated that task-related LC phasic activation occurs in relation to sensory stimuli, task-related cues and motor responses within cognitively demanding behavioral contexts such as decision making, attention, working memory and perceptual learning (Arnsten et al. 2012; Aston-Jones et al. 1994; Aston-Jones and Cohen 2005; Clayton et al. 2004; Glennon et al. 2019; Kalwani et al. 2014; Rajkowski et al. 2004; Robbins and Arnsten 2009; Sara 2009; Sara and Bouret 2012; Ghosh and Maunsell 2024). In most of this previous work, the appearance of a behaviorally relevant sensory stimulus was tightly tied to the motor output, creating a challenge in separating the role of LC activation in sensory and motor processes. Here, we sought to identify the distinct roles LC activation might play in sensory and motor components of perceptual decision making.

LC neurons provide the neuromodulator norepinephrine (NE) to the entire central nervous system via diffuse ascending axonal projections to most cortical and subcortical regions, as well as via descending projections to the spinal cord (Loughlin et al. 1986; Poe et al. 2020; Samuels and Szabadi 2008a). LC also receives projections from many brain regions; these inputs provide information about the current state of cognitive, sensory and autonomic processes throughout the nervous system (Poe et al. 2020; Samuels and Szabadi 2008b; Sara and Bouret 2012). There is now substantial evidence that different subsets of LC neurons selectively project to some brain regions but not others (Chandler 2016; Chandler et al. 2014, 2019; Kebschull et al. 2016; Uematsu et al. 2017; Poe et al. 2020; Chandler and Waterhouse 2012). Previous work suggests that subpopulations of LC neurons may have different behavioral functions associated with their differential projection targets (Hirschberg et al. 2017; Uematsu et al. 2017; Chandler 2016; Chandler et al. 2014, 2019). The LC-NE system is thus well posed to influence the brain wide sensory-motor processes underlying perceptual decision making behavior.

Previous work has shown that increased NE transmission can improve stimulus information processing, by improving the signal-to-noise ratio (SNR) of sensory stimulus evoked responses and sharpening sensory tuning curves, among other modulatory effects (Waterhouse and Navarra 2019). A recent study in macaques demonstrated a causal relationship between increased LC activation during attended visual stimuli and enhanced perceptual sensitivity, resulting in improved detection of changes in these attended visual stimuli (Ghosh and Maunsell 2024). Together, these findings suggest that the LC-NE system contributes to sensory processing during perceptual decision making. However, much about the role of LC in perceptual decision making remains to be clarified, including the timing and significance of LC activation in relation to an animal’s motor responses. Furthermore, how do LC neuron responses during cognitively demanding conditions relate to LC function during innate behavioral circumstances? One possibility is that there are subsets of LC neurons that activate in relation to any behavior that requires contextually-important sensorimotor processing, regardless of whether that behavior is instinctive or related to cognitive task demands (Bouret and Richmond 2009).

To clarify the significance of LC activation during perceptual decision making, we recorded from single LC neurons in two macaque monkeys while they performed a visual, two-alternative forced choice (2AFC), orientation change detection task in which perceptual decisions were reported by saccadic eye movements. Importantly, the trials in our task were structured such that the stimuli containing the sensory evidence for a decision were temporally distinct from the saccade target stimuli for reporting decisions. This task structure allowed us to thoroughly examine whether LC responses are important for sensory or motor aspects of the perceptual decision making process. Additionally, our task design allowed us to relate physiological LC activation to psychophysical performance, which is essential for assessing whether LC activation could influence perceptual ability.

We found that LC neurons in our recorded population did not respond to the sensory stimuli which contained the information used by monkeys to form their perceptual decisions. Furthermore, we did not observe any relationship between LC response magnitude and variability in perceptual accuracy of the monkeys. However, we consistently observed choice saccade-aligned activation across most LC neurons recorded in both monkeys. This motor related activation had both *pre*- and *post*-saccadic components, with pre-saccadic responses occurring in time to contribute to saccade preparation. Additionally, we found separate LC responses that were closely aligned with microsaccades which occurred after the monkey was presented with saccade target stimuli but was not yet cued to execute the motor response for reporting the decision. Overall, we identified a subgroup of LC neurons with motor-related responses whose function during perceptual decision making was clearly distinct from that of LC neurons contributing to improvements in sensory processing reported in other studies (Ghosh and Maunsell 2024; Martins and Froemke 2015). Our results provide novel evidence in support of a more universal role of LC activation in facilitating motor preparation and execution processes triggered by behaviorally-important sensory events.

## Methods

### Subjects

Two adult rhesus macaques (Macaca Mulatta) were used for this study, one female (Monkey DO) and the other male (Monkey WA). Before beginning behavioral training, we affixed a titanium headpost to the skull of each monkey in a sterile surgical procedure under isoflurane anesthesia for the purpose of immobilizing the head during experiments. Experimental procedures were approved by the Carnegie Mellon University Institutional Animal Care and Use Committee and were in compliance with the United States Public Health Service Guide for the Care and Use of Laboratory Animals.

### Electrophysiology

After initial training, each monkey was implanted with a recording chamber (Crist Instruments; Hagerstown, MD) that provided access to the locus coeruleus. The chamber was placed on the midline (ML 0mm) and tilted in the AP plane at an angle of 30 degrees from vertical with the center of the chamber aimed at a location 8 mm above inter-aural zero. Each recording session began by lowering a single tungsten microelectrode (FHC; initial impedance 0.5 - 1 megaohm) through a sharp trans-dural guide tube. In both monkeys, the long guide tube length allowed the electrode to travel straight and come out approximately 23 mm below the dural surface. Next, the electrode was carefully advanced in depth through brain tissue, first at larger increments of 30um and later in smaller increments of 5um, while monitoring the neural activity to assess response characteristics of encountered neurons. A custom-made microdrive controlled by custom-written MATLAB software was used to lower the electrode. From session to session electrode trajectories could be reproduced reliably through the use of a plastic grid (1 mm inter-hole spacing) inside the chamber..

We used several strategies to find and validate the location of LC in the chamber. In both monkeys, we searched for and recorded from LC units located to the left of the midline. We first confirmed the chamber grid locations of the Superior and Inferior Colliculi (SC, IC). With our chamber positioning, the caudal edge of the SC (confirmed by evoking >30° amplitude saccades with electrical microstimulation) was located a couple of grid holes anterior to grid holes whose trajectories hit LC. Electrical stimulation of caudal SC neurons located in a grid location approximately 2 mm left of the midline evoked large upward saccades (>30° amplitude, angled at +25°) while stimulation of SC neurons in a grid location ∼3mm left of the midline evoked downward saccades. We used the topography of SC neurons to estimate the mediolateral position of LC in the chamber grid at ∼2mm left of the midline. IC neurons, identified by their obvious response tuning to different sound frequencies, were located in a couple of grid holes posterior to caudal SC. We also located the trochlear decussation and the trigeminal nerve/nucleus (me5/Me5), which reside in close proximity to LC. We identified the trochlear decussation by observing characteristic ‘buzzing’, high firing rate, ramp-and-hold responses to saccades with downward and diagonal direction components. We identified the mesencephalic trigeminal (Me5) axonal tract by observing neural signals closely aligned with mouth movements made by the monkeys. Finally, with the knowledge of the positioning of the landmark brain regions described above, we searched for a cluster of neurons with response characteristics matching known LC response properties from previous studies. Potential LC units were checked for waveform shape, low firing rate, sensitivity to startling/loud noises, and sleep-wake transitions in response rates as reported by previous studies (Aston-Jones et al. 1994; Bouret and Richmond 2009; Kalwani et al. 2014). Typically, the electrode had to be advanced another 7-9mm past the guide tube exit, before reaching LC. In both monkeys, all LC units were found in 2 grid holes spaced 1mm apart and located 1-2mm to the left of midline (presumably left hemisphere LC units). In monkey DO, we performed an MRI which confirmed that the trajectory of the recording chamber encompassed LC and showed that the visible electrode trajectories led to the generally correct region in the pons (see Results, Figure 1A). In Monkey WA, we recorded in two putative LC sites before, during and after an intramuscular injection of the α2-noradrenergic receptor agonist Clonidine (see Results, Figure 1B).

**Figure 1.**
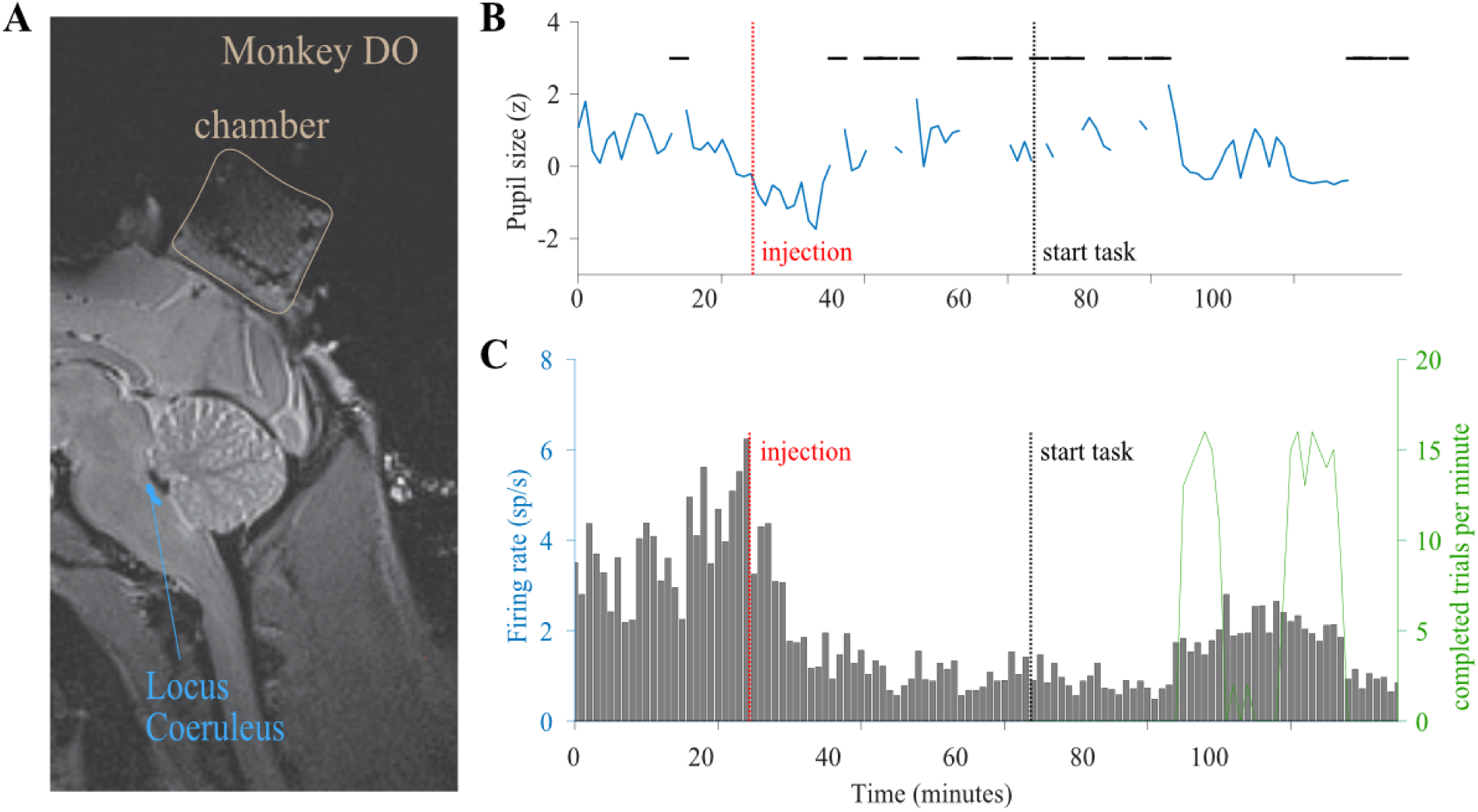
Confirming Locus Coeruleus (LC) Location. (A) A magnetic resonance image in the sagittal plane shows that the placement of the recording chamber (tan outline) allowed electrode trajectories down to the approximate location of LC (blue spot) in the brainstem of Monkey DO. Tracks are visible in the cortex marking damage from repeated guide tube and electrode trajectories down to LC. (B) In this session we performed a pharmacological test of a putative LC neuron with an intramuscular injection of an alpha-2 adrenergic receptor agonist (clonidine, 5ug/kg) in Monkey WA following a period of rest in which the task was disabled. (B) After the injection (red line), there was a decrease in pupil size (blue line) and increase in eye closing (horizontal black lines), and the animal became visibly drowsy. (C) After a brief increase in activity at the time of the injection (red line), there was a prolonged (∼ 1 hour) decrease in the responsivity of the LC neuron. After ∼1 hour, LC activity ramped back up just as the monkey was awake enough to engage in the change detection task (indicated by the # of completed trials per minute shown in green).

During data collection sessions, the response properties of each potential LC unit were tested before beginning the experiment. Recorded neural activity was band-pass filtered (0.3 – 7,500 Hz), digitized at 30 kHz, and amplified by a Grapevine system (Ripple, Salt Lake City, UT). Waveforms that crossed a threshold were recorded, saved and stored for offline classification. We manually set the threshold to allow recording of some noise and multi-unit activity in addition to the isolated single unit activity. We used custom spike-sorting software written in MATLAB (https://github.com/smithlabneuro/spikesort) to manually sort putative LC waveforms based on shape and inter-spike interval distributions (Kelly et al. 2007). For analyses, we included both well-isolated single units as well as multi-unit activity in cases where the waveforms of the involved units were practically indistinguishable in shape. We present results obtained from 23 separate sessions in Monkey WA during task version 1, 20 separate sessions in Monkey DO during task version 1, and 12 separate sessions in Monkey WA during task version 2.

### Behavior

During the experiments, stimuli were displayed on a 21” CRT monitor (resolution of 1024x768 pixels; refresh rate of 100 Hz), with a viewing distance of 36 cm. The visual stimuli were generated using custom software written in MATLAB (MathWorks, Natick, MA) and Psychophysics Toolbox extensions (Brainard 1997; Kleiner et al. 2007; Pelli 1997). We measured pupil diameter and eye position using an infrared eye tracking system (EyeLink 1000; SR Research, Ottawa, Ontario). On trials with correct behavioral outcomes, monkeys were rewarded with 2 drops of water delivered through a juice tube.

#### Behavioral Task

*Version 1.* The 2 monkeys performed the change detection task shown in Figure 2A. A trial began when the monkey fixated on a dot at the center of the screen. After a randomized time period (400-600 ms), two drifting grating stimuli appear on the right and left sides of the screen, but the animal had to maintain central fixation. In this first presentation, the grating stimuli were ‘samples’ and each had a particular orientation (either tilted left to 135° or right to 45°). These same sample grating stimuli were shown in each trial over the course of the session. The sample stimuli remained on screen for 350 ms, at which point they disappeared but the animal still maintained central fixation. After a randomized time period (200-400 ms), a ‘test’ grating stimulus appeared on the right OR the left side only, with a 0.5 probability of appearing on either side. This ‘test’ stimulus could have the same or a different orientation as the ‘sample’ stimulus that was first presented on that side of the screen. After 350 ms the second ‘test’ stimulus disappeared. Next, there was a variable delay period before the fixation point disappeared, and a green and red target circle appeared on the screen. The green target represented a “yes, change occurred” choice while the red target represented a “no change” choice. The disappearance of the central fixation point instructed the animal to report its choice with an eye movement to the green or red target. The position of the green and red targets between bottom and top locations was randomized session to session in monkey DO, and randomized trial to trial in monkey WA. Among the “change present” trials, the amplitude of the orientation change was selected randomly from 4 possible levels that were chosen based on the animal’s performance. “Change present” and “change absent” trials occurred with an equal likelihood. The animal received the same reward if he correctly indicated orientation change or no change, regardless of whether the stimulus was on the left or right side. Such task structure ensured that the animal’s behavioral strategy fluctuated naturally and was not biased by task conditions or selective signals such as spatial attention. Throughout each session, we used an inter-trial interval of 1 second. Monkey DO and Monkey WA completed 901 and 580 trials on average per session, respectively.

**Figure 2.**
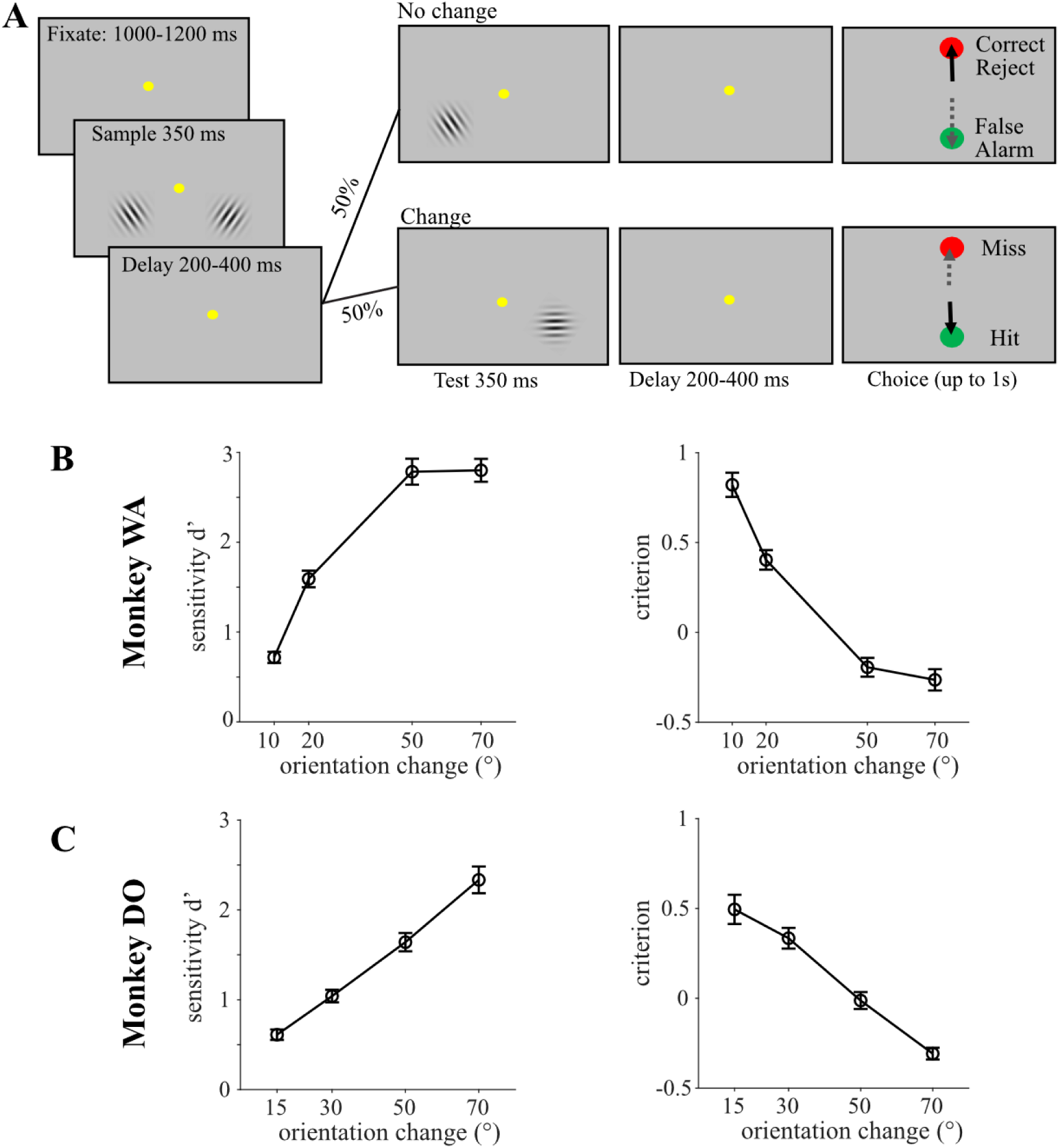
Behavioral task and performance. (A) An example trial in the perceptual decision making task. Two gratings were initially presented to balance the animal’s spatial attention. After a delay interval, monkeys reported whether the test grating either matched (no change) or did not match (change) the orientation of the sample grating at one of the two locations (randomly chosen on each trial) by making a saccade to 1 of the 2 colored choice targets (in this figure, green to indicate a change occurred and red to indicate no change occurred). Correct decisions were followed by juice reward within 1 second of the choice saccades. (B,C) Average psychometric functions across sessions for Monkey WA (B) and Monkey DO (C). The signal detection parameters sensitivity (d’) and criterion are plotted as a function of the difficulty (amplitude in °) of the orientation change.

#### Behavioral Task

*Version 2.* Monkey WA completed an additional 12 experimental sessions where he performed an alternate version of the change detection task (see Figure 6A). In this second version of the task, when the red and green targets appeared towards the end of each trial, the central fixation point remained on the screen for an additional randomized time period (350 – 550ms). Next, the fixation point disappeared (but the red and green targets remained on screen), cueing the monkey that it’s time to make the choice saccade. This version of the task was similar in all other ways to the first version described above.

### Data Analysis

#### LC activity

Baseline LC firing rates were computed in a 500 ms window during the initial fixation period in a trial, when no stimuli other than the fixation point were present on the screen. To compute LC firing rate responses to the ‘first stimulus grating’, or ‘second stimulus grating’, we used a 350 ms window aligned on the onset of the first or second orientation grating. We used a time window of +/- 100 ms centered on saccade onset for computing individual LC neuron firing rate responses to saccades indicating the monkeys’ choices on the task. We used a time window of (+/-100 ms) centered on choice target offset for computing LC firing rate responses during the trial event of all visual stimuli disappearing from the screen at the end of the trial. To assess the individual neuron responses to choice target onset, we computed LC firing rates in a window of 0-300 ms aligned on choice target onset.

For calculation of individual neuron peri-event time histograms (PETHs), we first counted spikes for each LC neuron in 20 ms bins within particular time windows during fixation (200 - 700 ms relative to fixation point onset), orientation grating stimuli (-200 to 500 ms relative to stimulus onset), the choice period (-200 to 300 ms relative to choice target onset), choice saccades (-300 to 200 ms relative to saccade onset), and choice target offset (-300 to 300 ms). To standardize a PETH for each LC neuron, we computed the mean and standard deviation of the distribution of each neuron’s baseline (fixation) responses across all completed trials in each session and then used these values for standardizing (z-scoring) each neuron’s responses to the different trial events. Thus, the assigned z-scores conveyed the number of standard deviations by which a neuron’s response to a trial event exceeded its baseline response. We compiled population PETHs across the standardized PETHs of individual neurons.

For the analyses depicted in Figure 6A and Figure 7C, for each individual neuron, we first separated the motor response into pre- and post-saccadic components, then tested whether the magnitudes of these pre- and post-saccadic responses were significantly higher than a pre-choice period baseline firing rate computed in a time window that just preceded the onset of the green and red choice targets (-350 to -250 ms) for Figure 5A and (-650 to -550 ms) for Figure 6C, both aligned on saccade onset). For the scatter plots depicted in Figures 5A and 6C, we subtracted the average pre-choice baseline firing rate from each individual neuron’s firing rates computed in time windows just before (-100ms to 0) and just after (0 to 100ms) a saccade. This baseline subtraction measure allowed us to separate, quantify and compare the magnitudes of the pre- and post- saccadic responses.

#### Microsaccade detection

We used a 2D-velocity based algorithm (Engbert and Kliegl 2003) to determine a velocity threshold that eye movements had to exceed to be classified as potential microsaccades. For each completed trial, we computed the velocity threshold as a multiple (4) of the standard deviation of the distribution of velocities across a time period starting at -300 ms relative to the onset of the first stimulus grating and ending at fixation point offset (the ‘go’ cue for making a larger saccade). We defined microsaccades as any eye movements with a velocity between this lower bound threshold and an upper bound of 100°/sec. We removed any eye movements with an amplitude of greater than 1°, or a duration of less than 6 ms. We visually inspected eye position traces (see Figure 8A) in some sessions to verify the validity of our microsaccade detection method.

#### Quantifying behavioral performance

In the orientation change detection task used in this study, monkeys could report Yes (green) or No (red) decisions about the presence or absence of a stimulus change on each trial. This task design is rooted in signal detection theory and yields 4 possible behavioral outcomes: correct (hit), correct reject, miss, and false alarm (FA). Hit and miss outcomes occurred when a change in stimulus orientation occurred and the monkey made a saccade to the green (‘yes’) or red (‘no’) choice target, respectively. Correct reject and false alarm outcomes occurred when there was no change in the stimulus orientation, and the monkey made a saccade to the red (‘no’) or green (‘yes’) choice target, respectively. We used these trial outcomes to quantify behavioral performance. Hit rate was calculated for each session as the number of hit trials divided by the total number of trials on which a stimulus change happened (hit + miss trials). False alarm rate was calculated as the number of false alarm trials divided by the total number of trials on which a stimulus change did not happen (false alarm + correct reject trials). Criterion (c) and sensitivity (d’) were computed using signal detection theory. Sensitivity was approximated as d’ = *invcdf*(Hit Rate) – *invcdf*(FA Rate), where *invcdf* is the inverse cumulative density function of the normal distribution. A larger d’ value indicates better sensitivity to stimulus change. Criterion was approximated as c = -0.5 *(*invcdf*(Hit Rate) + *invcdf*(FA Rate)). A criterion value of 0 indicates that a subject has no bias for reporting the absence or presence of change in a stimulus. When c < 0, the subject is biased towards reporting that there was a stimulus change (green, ‘yes’) and when c > 0, the subject is biased towards reporting that there was no stimulus change (red, ‘no’). For psychometric functions shown in Figure 2, we calculated d’ and criterion from Hit and FA rates associated with each amplitude of the orientation change.

#### Statistical significance tests

To assess whether the magnitude of each neuron’s response to a trial event (i.e., grating stimulus, choice targets, pre- and post- saccade) was significantly different from a baseline response, we used the paired-sample t-test (Figures 3, 5, 6). To compare population (non-standardized) LC firing rate responses across different amplitudes of orientation changes or across behavioral outcomes (Figure 4), we used a one-way ANOVA with Tukey-Kramer post hoc comparisons to test for significant differences between groups (at p<0.05). To assess whether each neuron’s response to choice microsaccades was significantly different from its response to ‘other’ microsaccades (Figure 7), we used a two-sample t-test (with the Welch’s correction for the assumption of equal variances) because we had unequal sample sizes.

**Figure 3.**
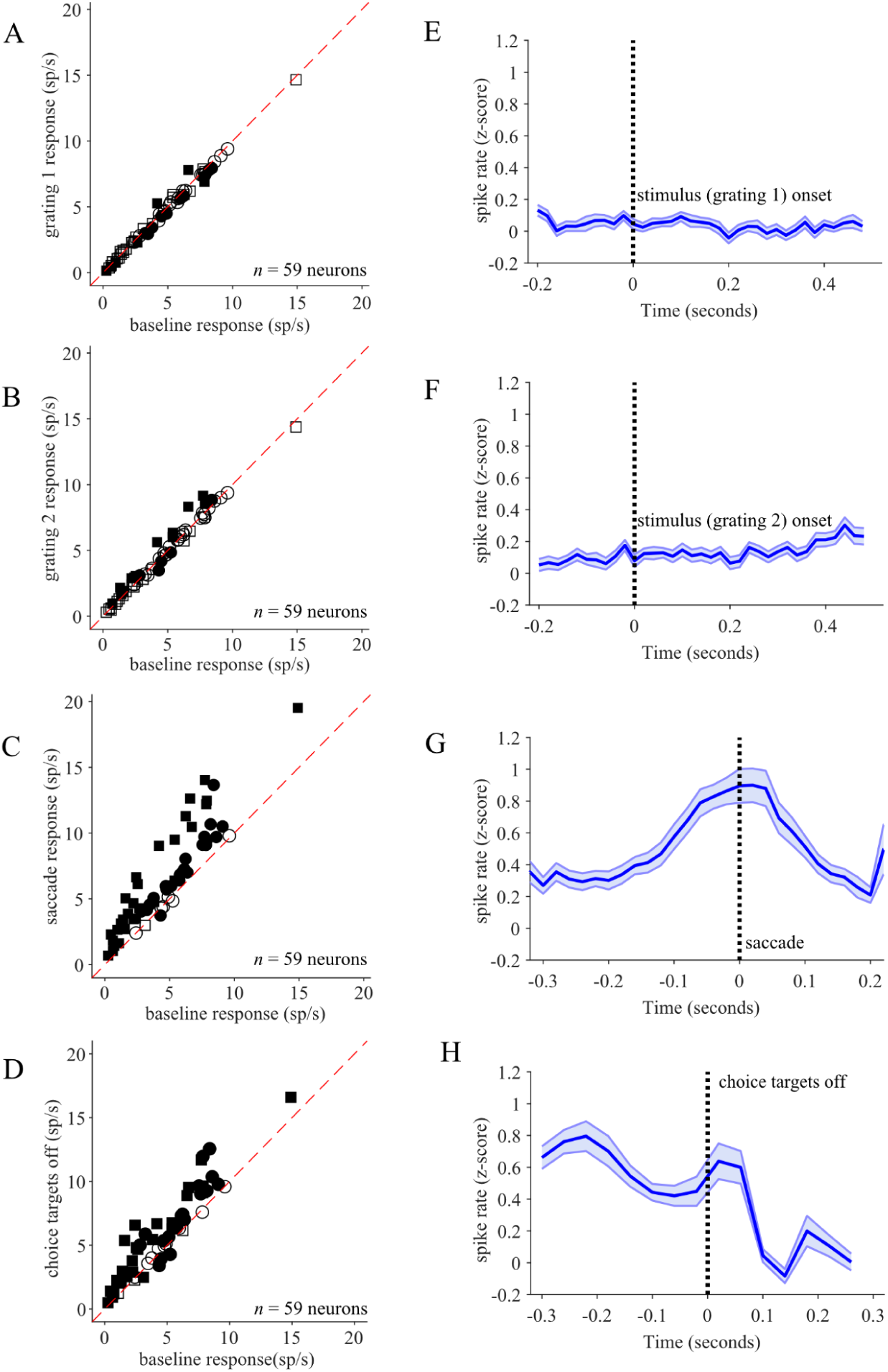
Task-related LC phasic activation. (A-D) Scatter plots depict the responses of all recorded individual LC neurons aligned to the various task periods: the first stimulus grating onset (A), the second stimulus grating onset (B) the saccade indicating the monkey’s choice (C), and the disappearance of choice targets (D) as compared to each neuron’s baseline response (x-axis). The squares (WA) and circles (DO) represent neurons recorded from the two different monkeys, and the filled points indicate LC responses significantly different from baseline (paired t-test, p < 0.05). The majority of the 59 recorded LC neurons showed significant elevation in activity around the time of the choice saccade. (E-H) Average PETHs across all standardized LC responses (blue bold line and shading represent mean +/- SEM across Z-scored PETHs of individual neurons) aligned on onset of first stimulus grating (E), onset of second stimulus grating (F) and choice saccade onset (G) and the disappearance of choice targets (H).

**Figure 4.**
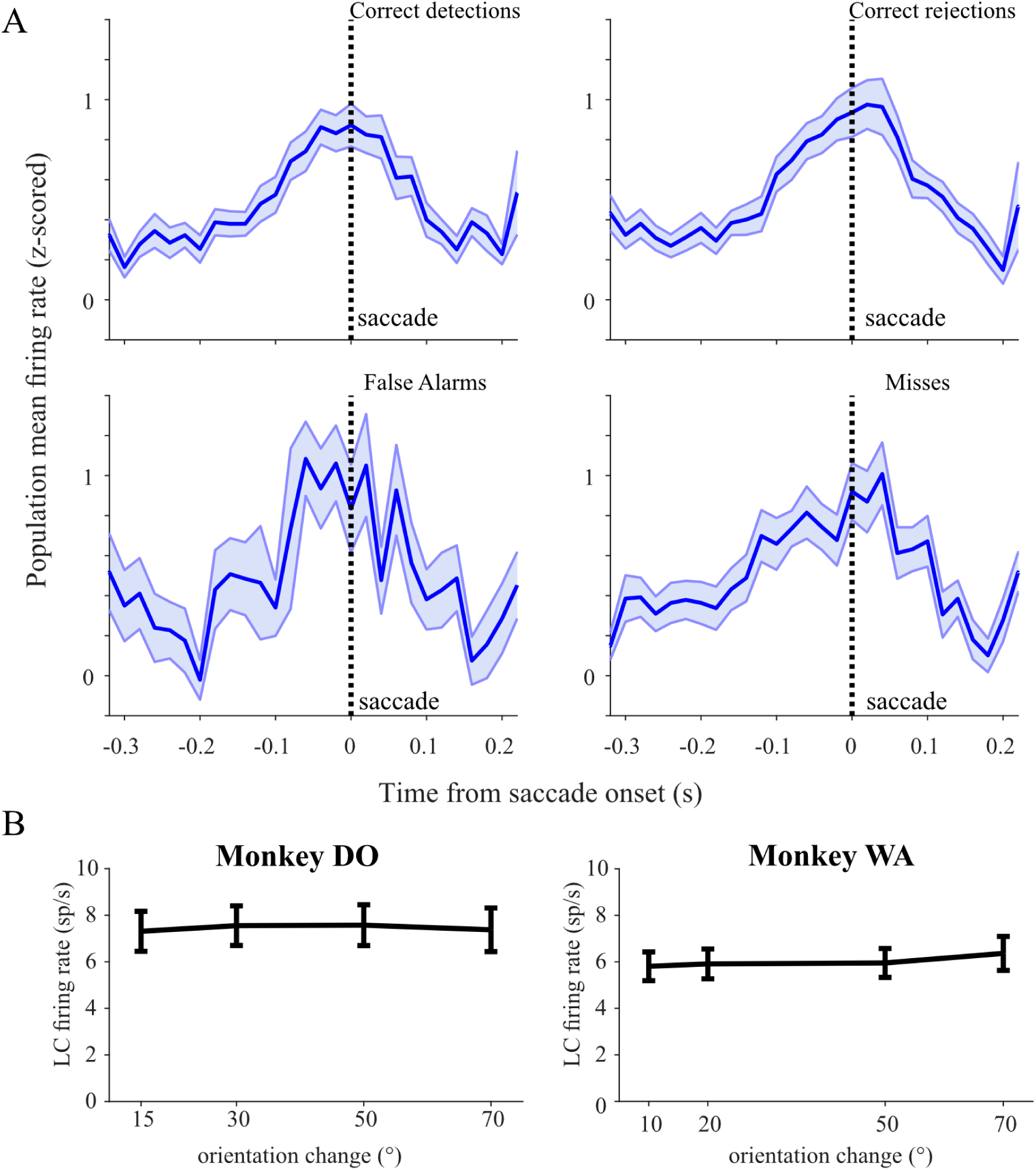
Peri-saccadic activation of LC neurons contingent on trial outcome. (A) In the same format as Figure 3G, average population PETHs are shown for the four possible trial outcomes. Because the overall performance across all trial types was above 50%, the numbers of trials in the false alarm and misses category were smaller and those PETHs are therefore noisier. (B) Saccade-aligned LC population average firing rates were equal across correct detections of each of the four orientation change difficulties (one-way ANOVA, post-hoc comparisons, all p>0.05 in Monkey WA and Monkey DO).

**Figure 5.**
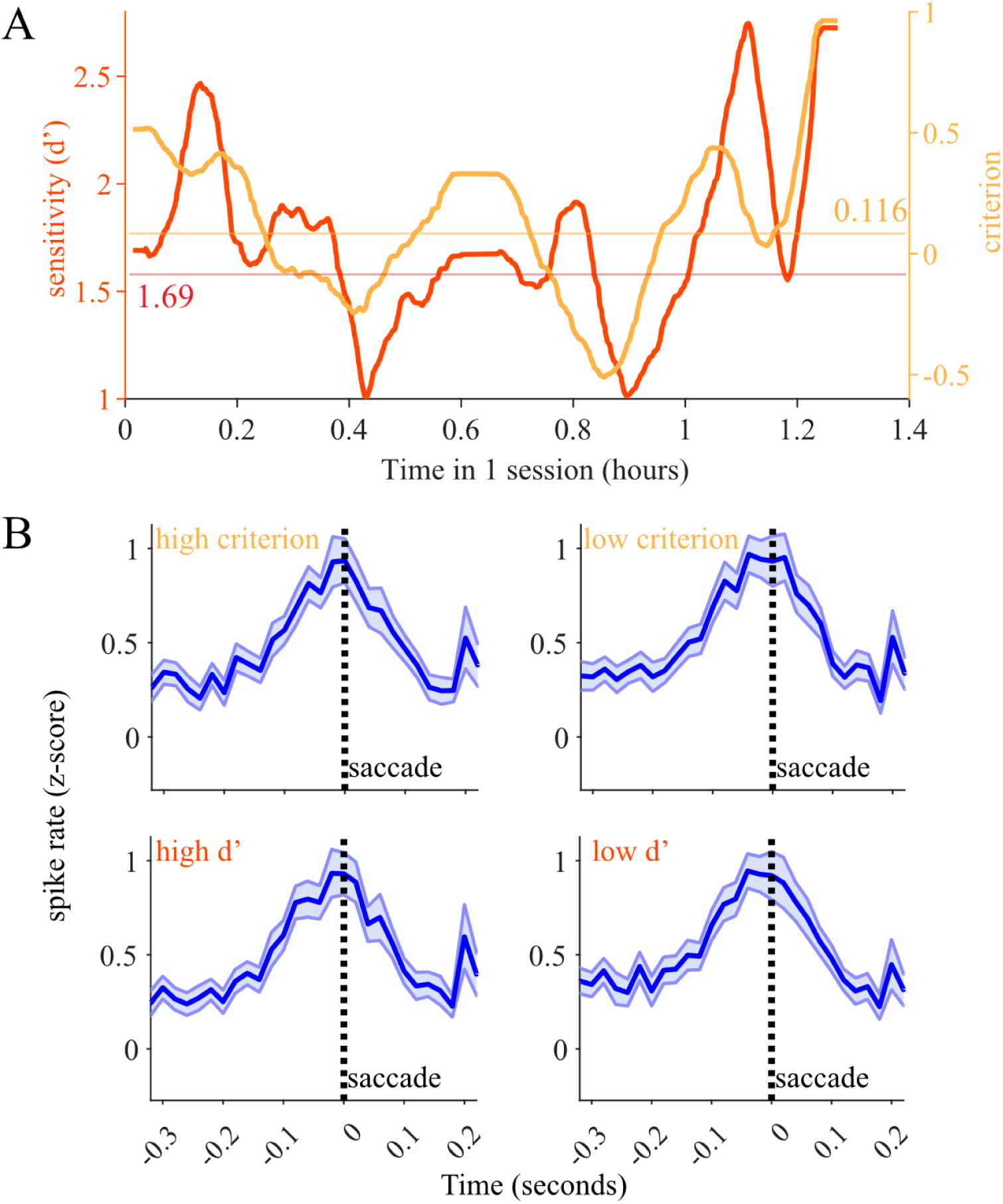
LC activation relative to slow changes in perceptual state. (A) We measured the perceptual sensitivity (d’, red) and decision making criterion (orange) in bins of 50 trials over the course of each session. We found that there were substantial fluctuations in behavioral performance over time. We labeled each trial as low or high criterion or sensitivity based on whether it occurred during a period of time in which those values were above or below the median throughout the session (indicated with horizontal lines). (B) There was no significant difference in saccade-aligned phasic activation between high and low criterion groups of trials (paired *t*-test, p = 0.41) Saccade-aligned phasic activation was also equal between low d’ vs high d’ groups of trials (paired *t*-test, p =0.90). Thus we found no evidence of changes in LC saccade-aligned responses relating to slow fluctuations in behavioral performance.

**Figure 6.**
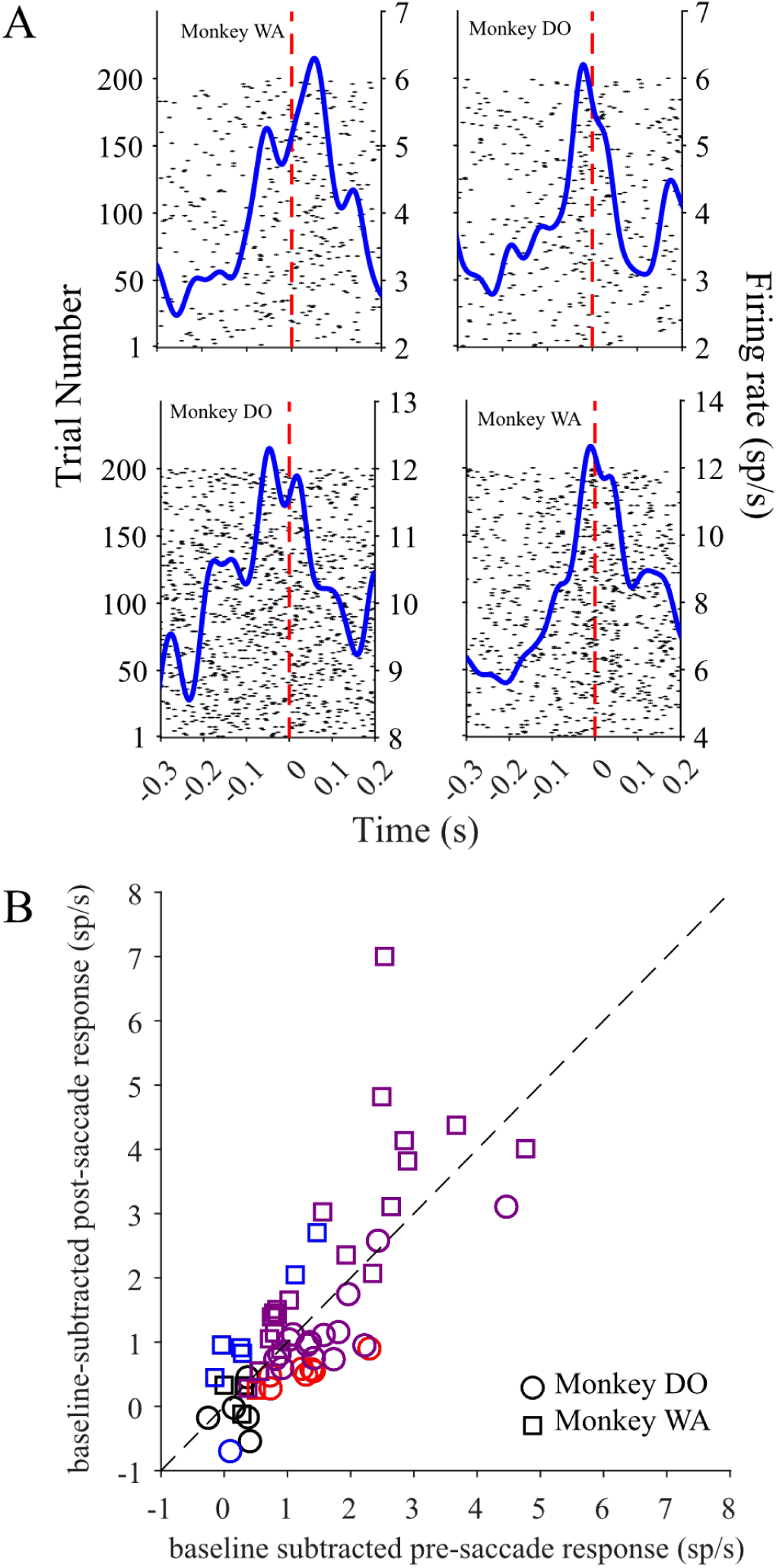
Time scale of peri-saccadic LC activity. (A) rasters and PETHs of four LC neurons around the time of the saccade demonstrate both pre- and post-saccadic activity. (B) Baseline-subtracted firing rates evaluated for each neuron in the 100 ms before (x-axis) and 100 ms after (y-axis) saccade onset revealed a relatively balanced distribution of timing of LC activity relative to the saccade. The red and blue symbols indicate neurons with significant pre-saccadic and post-saccadic activation, respectively. The purple symbols indicate neurons with significant elevation of activity in both time periods.

## Results

### Identification of LC Units

We recorded the activity of a total of 75 single LC neurons in two monkeys (n = 28 in Monkey DO during Task 1; n = 31 in Monkey WA during Task 1; n = 16 in Monkey WA during Task 2). Before beginning data collection experiments, we located LC by mapping the locations of the Superior Colliculus, Inferior Colliculus, trochlear decussation, and the tract of the trigeminal nerve (me5) in our recording chamber. These brainstem regions are typically considered good landmarks for finding LC because of their distinctive response characteristics and close proximity to the noradrenergic nucleus (see *Methods* for more details). Figure 1A shows an MRI image taken in Monkey DO after chamber implantation and several sessions of brain region mapping. The MRI confirmed that the positioning of the recording chamber allowed for electrode trajectories to pass through the estimated position of LC.

To further validate that we indeed correctly mapped the location of LC, on two separate days we administered an intramuscular injection of clonidine, an alpha-2-adrenergic receptor agonist, at a dose of 5ug/kg to Monkey WA while the electrode was resting in putative LC (Grant et al., 1988). We recorded the activity of putative LC units before, during and after the administration of clonidine. Consistent with previous reports of the effects of clonidine on LC activity (Joshi and Gold 2022; Joshi et al. 2016; Bouret and Richmond 2009; Kalwani et al. 2014), we observed that a few minutes after the injection, neural firing and pupil diameter decreased and the animal became noticeably drowsy, keeping his eyes closed for extended periods of time (Figure 1B). Approximately one hour after the injection LC activity ramped back up as the animal regained alertness and began to successfully complete trials in a visual perception task (Figure 1B).

In the beginning of each data collection session, we identified LC neurons based on several previously established characteristics: low spontaneous firing rates, broad and biphasic waveforms (cite), burst responses to startling sensory stimuli (e.g., knocking on door), and notable changes in responsivity as animals transitioned between periods of drowsiness and alert wakefulness.

### Task Performance

We trained two monkeys to perform a two-alternative forced choice (2AFC) orientation change detection task depicted in Figure 2. The monkeys reported the presence or absence of a change in the orientation of a drifting grating by making a saccade to either a green target (Yes) or a red target (No). We varied the difficulty of the orientation change to introduce some perceptual uncertainty as to whether each trial’s choice is correct or not. To analyze the monkeys’ behavioral performance, we applied signal detection theory, which is a method to statistically model the process of perceptual decision formation (Gold and Shadlen 2007; Macmillan and Creelman 2004; Shadlen and Kiani 2013). SDT is typically applied to binary (two-alternative) decisions, as in our task where the animal had to decide whether a change in orientation occurred or not. Reporting Yes (green) or No (red) for trials of the two possible stimulus conditions (change present or absent) provides 4 different behavioral outcomes on this task: hits, misses, correct rejections and false alarms (Figure 2A). There are two metrics that determine if a subject perceives or misses a stimulus change, these are 1) sensitivity (d’) and 2) criterion (c). Behavioral sensitivity is a measure of how well an ideal observer can separate the presence of signal from its absence while decision criterion conveys how strong the internal signal must be before the ideal observer decides to report a change.

Because changes in behavioral performance of the task could be due to changes in either criterion or sensitivity, we evaluated both parameters in both monkeys. We measured each monkey’s ability to detect orientation changes of different difficulties by calculating sensitivity (d’), and found that both monkeys had higher sensitivity to larger orientation changes, and lower sensitivity to smaller orientation changes (Figure 2B,C). Both monkeys had increased criterion at smaller change amplitudes and decreased criterion at larger change amplitudes (Figure 2B,C). These behavioral results are consistent with those reported by previous studies that utilized a similar task structure (Crapse et al. 2018). We conclude that the two monkeys utilized comparable decision making strategies to perform the task adequately, and as expected.

### Task-related LC Activation

Past studies of macaque locus coeruleus neurons have reported burst responses aligned with both sensory stimulus presentations and motor reports in decision making tasks (Clayton et al. 2004; Ghosh and Maunsell 2024; Kalwani et al. 2014; Rajkowski et al. 2004), but evidence for how these LC responses may contribute to perceptual decision making remains scarce. It is widely accepted that there are several stages to the perceptual decision making process, including sensory evidence evaluation, sensory-motor transformation, and motor preparation and execution. To more definitively connect the timing of changes in LC activity with specific elements of the perceptual decision formation process, we designed our task structure such that the stimuli containing the sensory evidence for a perceptual decision (the drifting gratings) were temporally distinct from the saccade target stimuli (green and red circles) for reporting choices. We recorded from single LC units while monkeys performed the task. The LC neuron response profiles, including response latencies and proportions of task-event responsive neurons, were similar for the two monkeys, allowing us to combine the two data sets for most of the analyses.

Increased LC activation occurred around four trial events: grating 1 and 2 onset, task-related motor action (saccades), and choice target offset at the end of the trial. We found that most recorded LC neurons exhibited elevated responses above baseline only during the choice period, right around the time of the saccade by which the monkeys reported their perceptual decision on each trial. The average baseline firing rate across all recorded neurons was 4.62 ± 2.93 spks/s. Figure 3C demonstrates that out of 59 recorded LC units, 52 neurons had responses aligned to choice saccade onset that were significantly higher than baseline responses during initial fixations at the beginning of trials (p < 0.05, paired sample t-test). In each monkey, a high proportion of LC neurons showed significant peri-saccadic activation during the choice period: 30/31 in Monkey WA and 22/28 in Monkey DO. We observed that a small subset of LC neurons (2/59) had significantly greater responses aligned to onset of the first stimulus grating, compared to baseline (p < 0.05, paired sample t-test; Figure 3A). Additionally, 15 of the 59 recorded LC units had responses to the onset of the stimulus 2 grating that were significantly elevated above baseline (p<0.05, paired sample t-test; Figure 3B). We confirmed that saccades to choice targets prompted the highest LC activation; the saccade-aligned responses were significantly greater than stimulus 2 aligned responses in 51/59 LC neurons (p < 0.05, paired sample t-test). We found that 43/59 LC neurons had significantly increased responses to choice target offset at the end of a trial (p < 0.05, paired sample t-test; Figure 3D).

Upon closer inspection of how individual neuron responses evolved over the course of a trial, we found that the recorded LC neurons had diverse response patterns within trials, activating in response to different combinations of trial events. The peri-event time histograms (PETHs) in Figure S3-1 highlight 5 individual neuron examples of the response heterogeneity we observed in the whole population. Some neurons spiked in response to the grating stimuli and the choice saccades within a trial (i.e. N33&37 in Figure S3-1), while others responded only to saccades and choice target offset during a choice period (i.e. N15 in Figure S3-1). In a small subset of neurons (5/59), spiking activity remained at baseline levels throughout the trial, decreasing immediately after the choice target offset at the end of a trial (i.e. N32 in Figure S3-1). Overall, fast transient activation was only observed in close temporal proximity with choice saccades and times of choice target offset.

We next examined the average response dynamics across the whole population of recorded LC units. To compile the population response, we first standardized the firing rate of every neuron to the mean and standard deviation of the distribution of baseline firing rates across all completed trials (see *Methods*). Figures 3E-H show the average peri-event time histograms (PETHs) of standardized LC responses aligned to the onsets of 4 important trial events: grating 1, grating 2, choice period saccade and choice target offset. Population activity remained at baseline levels during stimulus gratings 1 and 2 (Figure 3E,F). Consistent with our finding of increased firing by many individual LC neurons around choice saccade onset, we observed a prominent buildup of the population activity beginning approximately 100 ms before the saccade onset (Figure 3G). The increase in activity lasted through the saccade and gradually decreased during a 200 ms period following the saccade. Per task design, monkeys maintained gaze fixation at the location of the chosen red or green target for 200ms, at which point the choice targets disappeared from the screen and animals could look around freely. We observed a small increase in population activity within 50ms of choice target offset (Figure 3H). As confirmed by the population activity analysis, the most consistent response modulation across all the LC neurons recorded in this study was the motor-related activation before and after the choice saccade.

In one recent study, researchers recorded from LC neurons while macaques performed a visual orientation change detection task similar to the task described here, but with an added behavioral component of the animals selectively attending to one of two visual stimulus locations (Ghosh and Maunsell 2024). This study reported that a subset of LC neurons increased activation in response to an attended visual stimulus appearing in the hemifield contralateral to the site of the LC recording, and furthermore, the observed increase in LC activity was associated with enhanced perceptual sensitivity in task performance. Although our task design did not involve manipulating the animal’s spatial attention, other task parameters such as contralateral and ipsilateral presentations of drifting gratings were analogous. Therefore, we separately examined the activity of the subpopulation of 15 LC neurons with significantly elevated firing rates during the second grating to clarify whether spatially selective response modulation. This subset of LC neurons showed increased activity starting around the time of the first or second grating onset and lasting through the choice period (for example, see N33 or N37 in Figure S3-1). However, we did not observe a fast transient activation in response to the onset of the second grating, in the population activity PETH (Figure S3-2) or in the individual neuron responses (Figure S3-1). Furthermore, the LC neurons in this subgroup did not show a bias in response for ipsilateral versus contralateral presentations of the second grating (Figure S3-2).

In summary, the majority of LC neurons showed a fast transient activation around the time of motor action (the saccade) by which the animal reported a perceptual decision in an orientation change detection task. Notably, subsets of LC neurons responded to other trial events, such as the stimulus gratings, or the choice target offset in addition to activating during the choice saccade. This diversity of LC neuron responses reinforces the more recent theory of LC function which proposes that LC neurons activate as distinct ensembles with functions defined by anatomical connectivity rather than all LC neurons serving as a uniform source of global neuromodulation (Manger and Eschenko 2021; Poe et al. 2020; Totah et al. 2018). For the rest of this study, we focus on deciphering the behavioral significance and the timing of perisaccadic activation during the choice period, because LC neurons with significant motor responses comprise the largest proportion (88%) of recorded neurons in this study.

### Saccade-related LC responses are not modulated with changes in perceptual behavior

Previous studies have found that increasing extracellular norepinephrine concentration, via stimulating LC or by pharmacological means, can enhance responses of individual sensory neurons to sensory inputs. Additionally, computational models and a variety of studies in different species (rodents, monkeys, humans) have demonstrated that boosting NE transmission improves behavioral performance in perceptual tasks (Avery et al. 2013; Gelbard-Sagiv et al. 2018; Ghosh and Maunsell 2024; Guedj et al. 2019; Martins and Froemke 2015; Safaai et al. 2015; Servan-Schreiber et al. 1990). Despite numerous theories proposing a function for LC in perception and decision making, there is a currently a shortage of studies of LC activity in the context of demanding perceptual tasks that allow for the use of psychophysical measures to precisely quantify perceptual accuracy (Aston-Jones and Cohen 2005; Sara 2009; Waterhouse and Navarra 2019). To address this gap, we investigated whether the magnitude of saccade-aligned LC activation is modulated in relation to specific elements of perceptual decision making behavior including perceptual uncertainty, reaction times, perceptual sensitivity and criterion (as defined by SDT).

### LC Responses Are Not Modulated by Behavioral Outcome

To introduce some uncertainty into the decision making process, we varied the difficulty of the orientation changes that the monkeys had to detect. Accordingly, on some of the trials the monkeys were unsure of whether they saw a stimulus change or not, resulting in incorrect ‘false alarm’ change detections or ‘missed’ orientation changes. We considered whether LC phasic responses were modulated as a function of different behavioral outcomes of the monkeys’ choices. We grouped trials by the 4 possible behavioral outcomes on the task: correct detections, misses, correct rejects, and false alarms and compared LC population average responses between the conditions. Figure 4A shows that saccade-aligned LC phasic responses appear qualitatively similar in magnitude and temporal evolution across the 4 behavioral outcomes. Indeed, we found that the magnitude of LC activation was statistically indistinguishable across behavioral outcomes. This was true for both LC responses aligned to stimulus 2 grating as well as LC responses aligned on choice saccade onset (grating 2: one-way ANOVA, post-hoc comparisons, all p>0.05; saccade: one-way ANOVA, post-hoc comparisons, all p>0.05). Additionally, we verified that the subset of 15 grating 2-responsive LC neurons did not differ in responsivity across behavioral outcomes (grating 2-aligned: one-way ANOVA, post-hoc comparisons, all p>0.05; saccade-aligned: one-way ANOVA, post-hoc comparisons, all p>0.05). The similarity of LC responses across behavioral outcomes suggests that LC motor-related activation in this task is not related to any certainty or uncertainty the monkey has about its decision.

Additionally, we considered whether baseline LC firing rate (computed during the initial fixation period in each trial) differed between incorrect and correct behavioral outcome trials. In both monkeys, the average baseline firing rate across all recorded neurons was not significantly different between correct (hits and correct rejections) and incorrect (false alarms and misses) trials (Monkey WA: population mean firing rate on incorrect trials (3.9 sp/s) vs population mean firing rate on correct trials (3.9 sp/s), p = 0.87, paired t-test; Monkey DO: population mean firing rate on incorrect trials (2.7 sp/s) vs population mean firing rate on correct trials (2.8 sp/s), p = 0.81, paired t-test). Therefore, we conclude that the baseline (tonic) LC firing rate did not predict the behavioral outcome of the monkey’s perceptual decision process on each trial of the task.

### Relating Perisaccadic LC Activation to Perceptual Sensitivity, Criterion, and Behavioral Response Times

While performing the task, both monkeys were less accurate at correctly detecting smaller amplitude orientation changes in the drifting grating stimuli. Since the monkeys’ choices were based on the available sensory information in presented visual stimuli, NE-mediated changes in how these stimuli are processed by brain regions along the visual pathway could affect the animal’s behavioral performance. We hypothesized that increased LC activity (and accompanying changes in NE efflux across the brain) during our perceptual decision making task may function to improve an animal’s perceptual accuracy.

We first asked whether LC response modulation reflected differences in the monkeys’ behavioral sensitivity across change amplitudes. For instance, the magnitude of LC activation could increase during harder change detections, resulting in improved SNR of visual cortex neuron responses, one of the known effects following increased NE transmission. Alternatively, LC spiking rate might track the monkeys’ behavioral sensitivity, with higher firing rate corresponding to improved detection ability of larger amplitude changes. We compared LC population average firing rates across the four orientation change difficulties that occurred in each monkey’s task. Because the range of orientation change amplitudes used for Monkey WA differed from the range used for Monkey DO, for this analysis we considered LC neurons recorded in each animal as separate populations. In both monkeys, we found that saccade-aligned LC population average firing rates were equal across correct detections of each of the four orientation change difficulties (Figure 4B, one-way ANOVA, post-hoc comparisons, all p>0.05 in Monkey WA and Monkey DO). Grating 2-aligned population average responses were also equal across the difficulty conditions (one-way ANOVA, post-hoc comparisons, all p>0.05 in Monkey WA and Monkey DO). This result demonstrates that saccade-linked LC responses are not encoding the perceptual effort required to detect more subtle orientation changes, nor are they related to the monkey’s ability to detect stimulus changes of different difficulties. However, motor-aligned activation during our task could be related to other aspects of perceptual decision making.

We next considered whether the latency or magnitude of LC activation during the choice period is related to behavioral response times (RT). For each monkey, we grouped all completed trials within each session based on whether a trial’s RT was higher or lower than the median RT of the session. The mean ‘long’ and ‘short’ RTs were comparable between the two monkeys, so we pooled the data for the following LC response analyses (Monkey WA: 207 ms and 174 ms; Monkey DO: 236.1 ms and 169 ms). Across sessions and monkeys, the mean ‘short’ RT was 170.6 ms and the mean ‘long’ RT was 223.5 ms (Figure S-4). Upon a qualitative examination of LC activity aligned on the onset of the choice targets, we did not observe an obvious difference in the magnitude or latency of population average LC phasic responses between short vs long RT trials (Figure S-4). Quantitatively, there was no significant difference in LC firing rates aligned on the choice period onset when compared between long and short RT groups (paired t-test, p =0.89). Upon inspecting saccade-aligned PETHs, we observed that LC activity on shorter RT trials increased at an earlier latency compared to longer RT trials, about 150 ms before the saccade. Otherwise, the saccade-aligned LC phasic responses appeared nearly identical between the short and long RT trials. There was no significant difference in the magnitude of the saccade-aligned LC responses when compared between long and short RT groups (Figure S-4; paired t-test, p = 0.5).

The difference in response times observed across trials likely stems from trial-to-trial variability in decision processes. For instance, the RTs could be affected by the difficulty of the sensory information interpretation. Indeed, in each monkey, we found that the average response time for correct detections of easy orientation changes was significantly shorter compared to average response time for detections of hard orientation changes (Monkey WA: 184 ms vs 190.2 ms, paired t-test to compare means across sessions, p = 0.016; Monkey DO: 161.5 ms vs 195.3 ms, paired t-test, p = 1.2e-16). Additionally, we found that in both monkeys, average RTs on correct rewarded trials were slightly shorter than RTs on incorrect unrewarded trials (Monkey WA: 189 ms vs 194 ms, paired t-test to compare means across sessions, p = 0.021; Monkey DO: 197.8 ms vs 217 ms, paired t-test, p = 5.7e-18). However, as reported above we did not observe a significant difference in LC response magnitudes between trials of different behavioral outcomes or different change amplitudes (Figures 4,5). We conclude that although LC activation is closely associated with within-trial task relevant behavioral responses, there is no LC response modulation that could explain why behavioral response times are longer on some trials than others.

Several studies, including one from our lab, have shown that perceptual performance fluctuates over the course of a session (Cowley et al. 2020). One previous report suggested that changes in levels of LC spiking activity over time may track fluctuations in behavioral performance during a simple target detection task (Usher et al. 1999). Thus, we next considered whether trial-to-trial fluctuations in magnitude of LC saccade-related responses reflected changes in the monkeys’ perceptual decision making behavior over the course of a session. We measured detection sensitivity (d’) and criterion within windows of 50 trials, shifting the analysis window in 1 trial increments over the course of each session. This yielded an estimate of the animal’s perceptual state for each trial. Figure 5A demonstrates an example session where both sensitivity and criterion fluctuated over the time that Monkey WA performed the change detection task. We found similar behavioral variability over the course of most of the recording sessions in both monkeys. We then grouped all within-trial LC responses by whether they occurred in a period when sensitivity or criterion was lower or higher than the median for the session. This median-based grouping strategy allowed us to split trials into two equally sized data sets in each session.

Figure 5B shows saccade-aligned PETHs of LC responses during high and low d’ trials as well as high and low criterion trials, combined across sessions and monkeys. We found no difference in the average saccade – aligned LC population response between low d’ vs high d’ groups of trials (n=59 neurons in low and high d’ conditions, paired t-test to compare population means; aligned to saccade, p =0.90). Similarly, we found no difference in the average LC population response between low criterion vs high criterion groups of trials (paired t-test; aligned to saccade, p = 0.41). In summary, although we consistently observed LC activation in close temporal proximity to decision reports during our perceptual decision making task, these LC signals do not appear to encode anything related to how accurately the monkeys process and interpret the sensory evidence provided by the presented visual stimuli.

### Time course of saccade-related response

Evidence from previous studies diverges on whether LC activation occurs before or after motor action onset. In the context of behavioral tasks where monkeys are trained to ‘lever press’ or perform another limb movement to complete trials, LC neurons reliably activate just before motor action onset (Bornert and Bouret 2021; Bouret and Richmond 2009, 2015; Jahn et al. 2020; Rajkowski et al. 2004; Varazzani et al. 2015). In two recent studies, macaques engaged in decision making tasks by initiating saccades, and these saccadic eye movements were associated with fast transient post-saccadic LC activation (Ghosh and Maunsell 2024; Kalwani et al. 2014). It is important to more precisely characterize the temporal course of the saccade related LC response within our perceptual decision making task, as the behavioral implications differ depending on whether LC activation begins before or after the motor action. The time course of LC population activity in our study suggests that most LC neurons in our data set activate before the saccade initiation, and this activation lasts until after the saccade is completed (Figure 3G). Upon inspecting individual neuron PSTHs, we found many instances of both *pre*- and *post*-saccadic peaks in response (i.e. the units depicted Figure 6A). We next quantified whether all individual neurons had a comparable activation throughout the saccade, or whether subgroups of neurons had distinct *pre*- or *post*- saccadic responses. We separated the motor response during the choice period into pre- and post-saccadic components by computing firing rate in two time windows - one immediately preceding and the second immediately following the saccade ([-100ms 0ms] and [0ms 100ms] respectively). We computed a baseline-subtracted measure of pre- and post-saccadic firing rates shown in Figure 6B, in order to clarify whether any detected pre-saccadic response was greater than merely a recent baseline rate increase that sometimes occurred towards the end of a trial (see Methods for details).

Increased *pre*-saccadic and *post*-saccadic activation occurred in 36/59 recorded LC neurons (p < 0.05, paired sample t-test, purple markers in Figure 6B). The example raster plots and PETHs in Figure 6A depict increased spiking activity within a +/- 100 ms time window aligned on the choice saccade, with some neurons showing distinct *pre*- and *post*-saccadic peaks in response. 44/59 neurons showed significantly increased *pre*-saccadic activation compared to the pre-choice period baseline activity (p < 0.05, paired sample t-test, red markers in Figure 6B). 43/59 neurons showed *post*-saccadic activation that was significantly higher than the baseline activity (p < 0.05, paired sample t-test, blue markers in Figure 6B). A small number of neurons (9/59) did not have appreciably elevated activation in either the pre-saccadic or post-saccadic time windows black markers in Figure 6B). Comparing across the two monkeys, similar proportions of LC neurons had significantly increased responses before the saccade (monkey D: n = 22/28; monkey W: n = 22/31) and after the saccade (monkey D: n = 16/28 units; monkey W: n = 27/31 units). Together, these results show that both *pre*- and *post*-saccadic responses were prevalent across the population of recorded LC neurons. The majority of LC units (75%) show motor-related responses that begin in time to influence the upcoming saccade, implying that LC activation contributes to motor preparation in the context of perceptual decision making.

### LC Activation during Choice Period: sensory, motor or both?

Overall, the results presented so far suggest that LC activation observed during the choice period of the task is more related to preparing and reporting a behaviorally important decision, but not to any decision process elements related to perceiving and interpreting the sensory information. However, we still have not answered the question of whether the observed LC responses during the choice period are sensory, motor, or both in nature. That is, is the activation during the choice period a combination of a sensory response signaling the onset of choice targets and a motor-related activation signaling an impending behavioral act, the saccade? To address this question, we collected an additional data set in Monkey WA while he performed a slightly altered version of the change detection task (see Methods). Our main goal with the task change was to separate the timing of choice target onset from the ‘go cue’ signaled by fixation point offset (Figure 7A).

**Figure 7.**
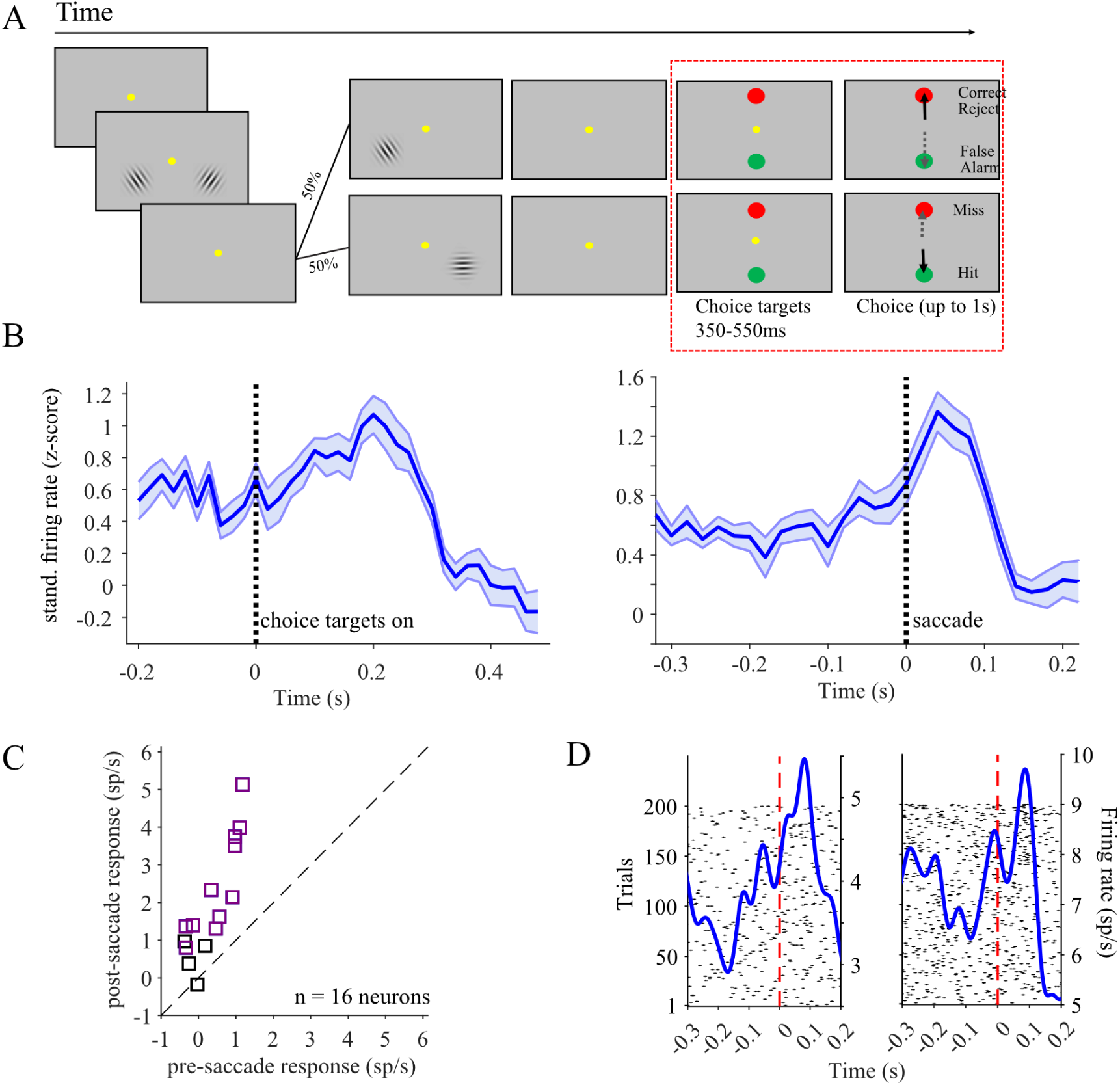
Separation of choice and saccade intervals in a control task. (A) We designed an alternative version of the task performed only by Monkey WA in which the choice targets appeared earlier before the go cue. The red dashed line shows the period of time in this control task which differs from the main task shown in Figure 2A. We recorded and analyzed the activity of 16 LC neurons during this task. (B) A population average PETH aligned to the choice target (left) showed a peak response around 200 ms after target appearance. A PETH aligned to the saccade onset (right) showed a peak roughly 50 ms after saccade onset. (C) Individual neurons in these sessions tended to have stronger post-saccadic than pre-saccadic responses, though a majority (12/16) showed significant responses above baseline both before and after the saccade. (D) Two individual neuronal PETHs and rasters indicate that the peri-saccadic response begins before saccade initiation and peaks after.

While Monkey WA performed the task, most recorded LC neurons showed distinct responses for the onset of choice targets and the saccade. Figure 7B demonstrates average target onset aligned and saccade aligned PETHs assembled across the activity of 16 LC units recorded during the altered version of the task. We found that 15/16 LC neurons had choice target onset aligned responses that were significantly higher than baseline responses (p < 0.05, paired sample t-test). Furthermore, the same 15 LC neurons also had responses aligned to choice saccade onset that were significantly higher than baseline responses (p < 0.05, paired sample t-test). We found that 6 out of the 15 neurons had saccade-aligned activation that was higher than target onset – aligned activity (p<0.05, paired sample t-test). Notably, pre-saccadic activation persisted in this version of the task: we detected significant *pre*- and *post*-saccadic activation in 12 of the 16 recorded neurons (p<0.05, paired sample t-test; Figure 7C). The example neuron raster and PETH plots shown in Figure 7D demonstrate that the increase in saccade-related activation begins before saccade initiation, and this pre-saccadic activation is separate from the target-onset aligned response.

Because the monkey could start planning the saccade as soon as the choice targets appeared on the screen, we hypothesized that the observed target onset – aligned LC responses could be related to motor planning rather than the sensory aspects of the choice targets. To address this possibility, we first grouped trials by the amount of variance in the monkey’s eye position (fixation variance) throughout the time period between choice target onset and the ‘go’ cue onset. We found that choice target onset – aligned LC activity was higher in trials with higher fixation variance compared to trials in which the monkey’s eye position was more stable during fixation (not shown). Next, we more closely examined traces of horizontal and vertical eye position in each trial in order to detect any small fixational eye movements that the monkey might have made (see Methods). Although monkey WA maintained fixation throughout each completed trial, we observed that he occasionally made microsaccades which are small (<1° amplitude), rapid, typically involuntary deflections in eye position (Figure 8A). We grouped all detected microsaccades in a session by the time at which they occurred in a trial. This allowed us to examine whether there was a significant increase in LC activity when aligned on microsaccade onset, and also whether the magnitude of LC activation varied depending on the timing of the microsaccade within a trial. Figure 8B demonstrates that in monkey WA, LC activation was tightly linked with the onset of ‘choice period’ microsaccades, but not microsaccades that occurred during ‘other’ time periods in the trial (i.e., grating stimuli, initial fixation). We found that 15 of the 16 LC neurons had ‘choice period’ microsaccade aligned responses that were significantly increased compared to responses aligned on all ‘other’ microsaccades (unpaired t-test, p<0.05). Although the trial structure during the choice period differed between the two task versions, it was identical during the time in a trial before choice period onset. This allowed us to check whether LC activity aligned on ‘other’ microsaccades was similar across the two monkeys. As observed in Monkey WA, there was no increase in LC activity aligned on microsaccades that occurred outside of the choice period in Monkey DO (Figure 8C).

**Figure 8.**
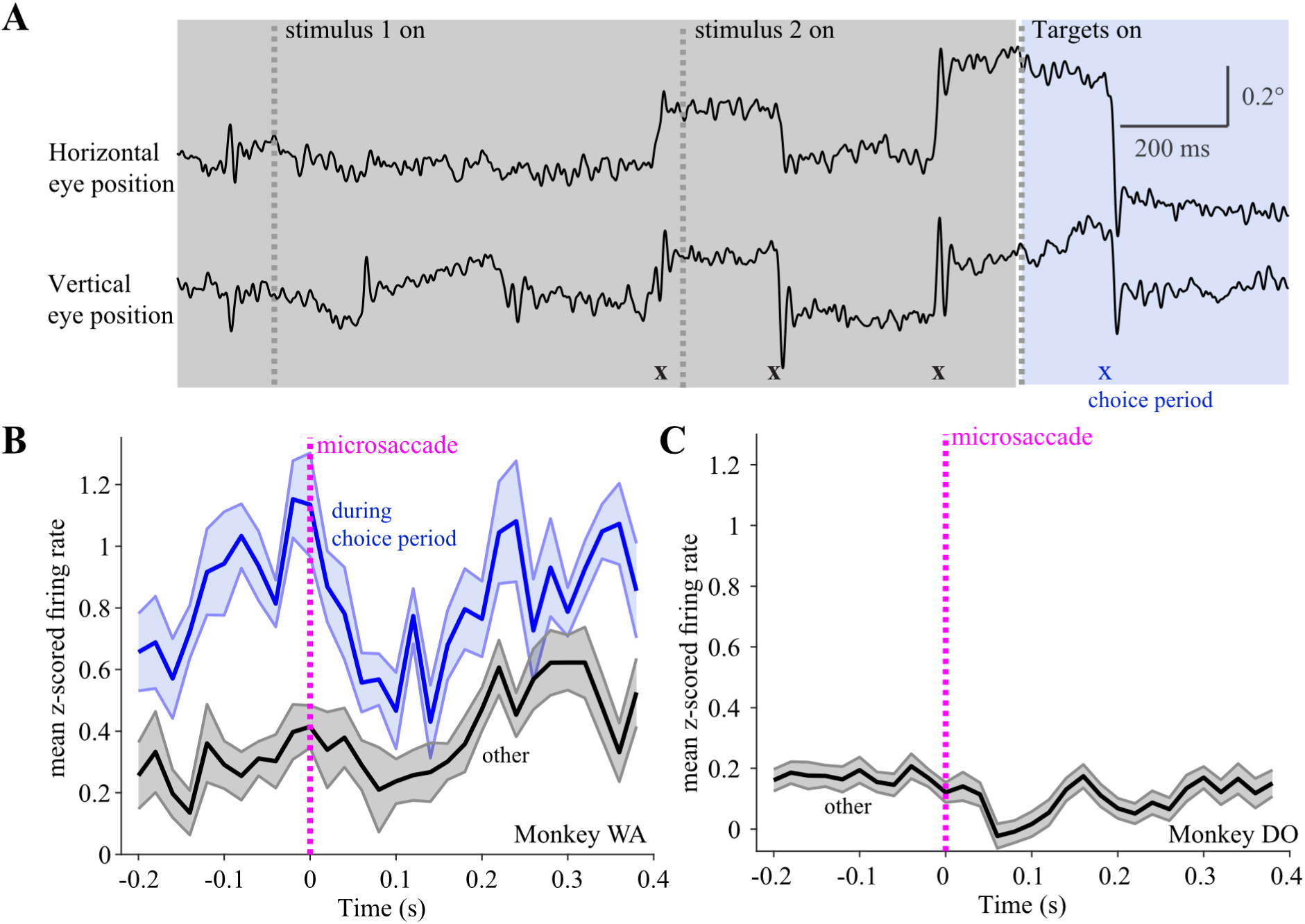
Relationship of LC activity to microsaccades. (A) During the fixation period of the task, animals made occasional microsaccades within the fixation window. These were detectable in individual trials by characteristic increases in radial velocity (indicated with X’s). We analyzed 16 LC neuron responses aligned to the onset of microsaccades (vertical pink dashed line) within trials of the control variant of the task (Figure 7) for Monkey WA (B) and 28 LC neuron responses within trials of the original task (Figure 2) for Monkey DO (C). For monkey WA (B), 15/16 LC neurons had ‘choice period’ microsaccade aligned firing rates (blue) that were significantly increased compared to their responses aligned on all ‘other’ microsaccades (black) that occurred within the trials but outside of the choice period (unpaired t-test, p <0.05). In addition, we confirmed that there was no increase in LC responses aligned on microsaccades that occurred outside of the choice period for monkey DO (C, black line).

## Discussion

We measured the responses of individual LC neurons while rhesus macaques engaged in a demanding perceptual decision making task that required them to detect changes in visual stimuli and report their decisions by making saccadic eye movements. Our findings provide novel insights into how transient, burst-like changes in LC activation within trials of the task relate to sensory and motor aspects of perceptual decision making. Although we consistently observed increased LC activation around the time of the saccade by which the animal reported a perceptual decision, we did not find any evidence for modulation of transient LC responses in relation to sensory information processing or variability in perceptual decision making behavior. Upon a closer examination of the time course of the perisaccadic responses, we established that a majority of the recorded LC neurons had both *pre*- and *post*-saccadic activation, with many pre-saccadic responses commencing in time to contribute to saccade preparation. In addition to choice saccade-aligned activation, we found separate LC responses that were closely aligned with microsaccades which occurred after the monkey was presented with saccade target stimuli but was not yet cued to execute the motor response for reporting the decision. To the best of our knowledge, microsaccade-aligned LC responses and pre-saccadic LC activation in the context of perceptual decision making have not previously been reported. Below, we relate our findings to existing work on the function of LC responses in decision making tasks and discuss how LC motor-related responses may function to orient and prepare an organism for a contextually appropriate and important behavioral response.

### Significance of LC Activation for Sensory Perception and Decision Making

In one of the earliest studies of LC activation during decision making, physiological LC activity was recorded while monkeys evaluated the directional orientation of presented visual stimuli (Clayton et al., 2004). The authors found that transient, burst-like (phasic) LC responses were tightly linked with the variable timing of hand movements by which the monkeys reported their decisions trial to trial. Based on these findings, the authors proposed that motor-aligned LC responses facilitate the execution of a behavioral response following the commitment to a particular decision. However, this and many other previous studies of LC activity during decision making (Aston-Jones et al. 1994; Clayton et al. 2004; Rajkowski et al. 2004; Usher et al. 1999) utilized a task structure that did not allow for a conclusive assessment as to whether the observed LC phasic activation contributes to both sensory information processing and motor processing components of a decision process. To address this point, we structured trials in our task such that the stimuli containing the sensory evidence for a decision were temporally distinct from the saccade target stimuli for reporting choices. Thus, we determined that the LC neurons in our recorded population did not respond in time to modulate the sensory processing and interpretation of the presented orientation grating stimuli within the same trial. Our results clearly show that most of the recorded LC neurons in both monkeys activated in close temporal alignment with eye movements during a specified decision reporting period, such that increased activation began in time to contribute to the upcoming saccade.

Our results do not preclude the possibility that changes in LC activity patterns can reflect arousal-related changes in behavioral performance in some behavioral contexts. A series of previous findings (Aston-Jones et al. 1994; Usher et al. 1999) formed the base for the prominent theory that transitions in LC activity patterns adaptively regulate behavior. This theory proposes that intermediate baseline LC activity and increased phasic activation serve to optimize performance on a current task, while elevated baseline activity and decreased phasic activation correspond to poor task performance, promoting increased distraction and exploration of other behavioral options (Aston-Jones and Cohen 2005). However, in our data, we did not find a difference in average LC baseline firing rates (computed during initial fixations within trials) when we compared responsivity between correct and incorrect behavioral outcome trials. We also considered whether LC phasic activation co-varied with behavioral performance over the course of an experimental session, but did not find a relationship between fluctuations in LC activation and changes in perceptual sensitivity and criterion, both measures of behavioral performance in a 2-AFC task such as ours (as shown in Figure 5). A few previous studies also failed to observe a relationship between baseline (tonic) LC firing rates and various metrics of behavioral performance (Kalwani et al. 2014; Rajkowski et al. 2004). It has been suggested that more extreme changes in arousal are required for the larger fluctuations in LC tonic activity that could influence an animal’s behavior (Aston-Jones and Cohen 2005). In contrast, LC activity appears to span some optimal range within trials of more perceptually challenging tasks such as ours, perhaps because considerable concentration is required for performing the task well enough to earn some reward.

Many previous studies have focused on manipulating either Locus Coeruleus activity or cortical NE concentration to elucidate the cellular and circuit mechanisms by which the LC-NE system might influence behavior. Increased NE transmission has been shown to improve the signal- to-noise ratio (SNR) of sensory stimulus evoked responses, alter sensory tuning curves, lower thresholds for sensory-evoked responses (gating), and enhance spike synchrony (Waterhouse and Navarra 2019). Together these previous findings prompted the prevalent theory that the LC-NE mediated improvements to sensory signal processing can alter perceptual behavior (Sara 2009; Waterhouse and Navarra 2019). A few studies have rigorously tested this theory by showing that changes in LC activity are linked with simultaneously recorded cortical sensory neuron responses in awake rodents engaged in a perceptual task (Martins and Froemke 2015; Yang et al. 2021). In rhesus macaques, the most conclusive evidence comes from a recent study that demonstrated a causal relationship between increased activation of a subgroup of LC neurons in response to an attended visual stimulus, and improved behavioral performance on a perceptual decision making task where animals were required to detect and report changes in grating orientation, similar to the task in this study (Ghosh and Maunsell 2024).

Based on these previous findings, we hypothesized that activation of LC neurons in trials of our change detection task would be linked with improved perceptual performance on the task. However, we found that most of our recorded population had motor-, but not grating stimulus-, aligned responses which did not signal changes in a subject’s perceptual accuracy. Our findings are consistent with a previous report that found no relationship between saccade-related LC activation and monkeys’ perceptual sensitivity to changes in attended visual stimuli (Ghosh and Maunsell 2024). Our results extend previous work on the contributions of LC to perceptual decision making, by showing that not all LC neurons are involved with enhancing sensory processing. Instead, we provide compelling new evidence of a subgroup of LC neurons with purely motor related function – which likely operates as such not only during perceptual decision making, but during other cognitively demanding and instinctive behaviors as well.

### Motor-related LC Activation Prepares an Animal for Important Behavioral Responses

Initial studies of the LC-NE system showed that unconditioned novel, salient or noxious stimuli of all sensory modalities evoke phasic responses, prompting hypotheses about the role of LC phasic activation in facilitating the processing of behaviorally important sensory information (Aston-Jones and Bloom 1981; Berridge and Waterhouse 2003; Foote et al. 1980). More recent studies have shown that some LC responses in various contexts including decision making are closely linked to the animal’s behavioral response (Bouret and Sara 2004; Clayton et al. 2004; Rajkowski et al. 2004). Furthermore, a number of recent studies have shown that LC neurons can have separable phasic responses to task-related conditioned sensory stimuli and behavioral action onset (Bouret and Richmond 2009; Bouret and Sara 2004; Kalwani et al. 2014; Varazzani et al. 2015). Similarly, in the second version of our task, we first observed an increased firing rate about 100 ms after the onset of the choice targets, followed by another transient activation around the onset of a saccade to one of the choice targets (Figure 7). It is noteworthy that in our task, some LC neurons selectively responded to the choice targets, but not to the stimulus gratings. That the LC neurons are able to respond differentially to 2 sensory stimuli containing distinct but behaviorally important information, suggests that LC phasic responses are tuned to a specific stimulus meaning. This is in line with previous studies showing that LC phasic activation updates following reversals in task contingencies that change the meaning of sensory stimuli (Aston-Jones et al. 1997; Dalley et al. 2001). Based on both previous observations and our results, it can be said that all the sensory stimuli which evoke responses from the motor-responding subgroup of LC neurons share the common characteristic of alerting or cueing the animal that a behavioral response may be necessary very soon in time.

What could be the function of these sensory and motor aligned LC responses that are so precisely timed to occur in very specific behavioral contexts? The timing of the spiking bursts could be merely correlated with important sensory cues and motor events because LC receives diverse afferent inputs that transmit contextually important information for the task at hand (Poe et al. 2020; Sara 2009; Sara and Bouret 2012). However, our results show that LC activation preceded the onset of microsaccades and saccades during the choice period, suggesting that LC activation occurs in time to modulate the preparation of these eye movements. Interestingly, previous work diverges on whether LC motor-related activation begins before or after the motor response. In experiments that designate a limb movement (i.e. arm lever press) as the task related behavioral response, LC activation precedes the motor action (Bouret and Richmond 2009, 2015; Clayton et al. 2004; Varazzani et al. 2015; Yang et al. 2021). However, LC activation was reported to occur after task related saccadic eye movements in the two studies in which saccade direction signified the behavioral response (Ghosh and Maunsell 2024; Kalwani et al. 2014). The timing of motor-related LC activation most likely varies depending on the behavioral context, as well as the anatomical location and connectivity profile of the LC neuron being recorded. These factors together likely also determine how LC motor related activation contributes to an animal’s behavior response. Below, we discuss how our findings provide new evidence in support of the theory that LC transient bursts facilitate processing in brain regions involved in motor output and autonomic activation, thus preparing the animal for an appropriate motor response to a behaviorally significant stimulus, whether that’s in a task setting or in the natural environment (Bouret and Sara 2005; Sara and Bouret 2012).

We found that in the altered, second version of our task (Figures 7,8) monkeys tended to make small fixational eye movements (microsaccades) while they waited for the offset of the fixation point which served as the ‘go’ cue for executing a larger saccade for reporting the decision. Importantly, during that time period, LC phasic activation was more tightly linked to the onset of the microsaccade than the onset of the choice targets (Figure 8). An important novel finding of our study is that, at least in the context of our task, the cue-stimulus aligned LC phasic activation did not reflect the sensory aspects of the choice targets, but rather signaled a motor process that was likely triggered by the onset of the choice targets. Although the visual function of microsaccades has been debated for decades, the current understanding is that microsaccades and saccades are the same type of eye movement, generated by a common oculomotor pathway (Martinez-Conde et al. 2013). For instance, there is now substantial evidence that neurons throughout the Superior Colliculus represent all saccade directions and amplitudes, with the <1deg amplitudes typical of microsaccades encoded by neurons in the rostral pole of SC (Hafed et al. 2009; Munoz and Wurtz 1993). Just as saccades have multiple roles in vision, microsaccades may also serve several important functions, including correcting fixation errors (Engbert and Kliegl 2004), restoring fading vision during fixation (McCamy et al. 2012), sampling the available information in the visual environment (Martinez-Conde et al. 2013), and overtly signaling shifts in covert attention (Engbert and Kliegl 2003; Hafed and Clark 2002; Yu et al. 2022; Willett and Mayo 2023).

It is possible that in our task LC activation facilitated microsaccades during the choice period to help the monkey covertly orient to the locations and colors of the two choice target stimuli. Of note, we only observed significant LC activation around microsaccades during the choice period, but not microsaccades occurring during other times in the trial (Figure 8). Such selective timing implies that LC phasic responses and the corresponding NE release in target regions function to optimize motor preparation for an upcoming important behavioral response, which in our task was a saccade to report a decision. Our interpretation is consistent with previous anatomical tracing studies which have shown that within the primate visual system, LC projections predominately target the prefrontal cortex and tecto-pulvinar structures, while projections to the visual and inferotemporal cortex are comparatively less dense (Arnsten 1998; Lewis and Morrison 1989; Morrison and Foote 1986; Porrino and Goldman-Rakic 1982). Others have noted that such a projection pattern supports a role for the LC-NE system in preferentially regulating activity in brain regions with visuomotor and spatial analysis functions, such as the superior colliculus (Berridge and Waterhouse 2003; Morrison and Foote 1986).

Interestingly, recent studies provide compelling evidence that action-onset aligned LC phasic activation signals the amount of physical effort required for a task-relevant important behavioral response (Bornert and Bouret 2021; Varazzani et al. 2015). These findings support the theory that motor-related LC phasic activation promotes mobilization of physical (i.e., muscles) and autonomic (i.e., respiratory, cardiac systems) resources to help complete physically effortful actions. Our results are consistent with such an interpretation and extend previous understanding of LC motor-related activity in two important ways. Firstly, it can be said that behavioral responses in the above-mentioned studies were by nature volitional, as monkeys had to exert a specific understood amount of physical force for completing trials. Because microsaccades are typically involuntary, our finding of LC activation around microsaccades provides evidence that LC neurons respond to both voluntary and involuntary motor outputs. Secondly, although the physical effort required for behavioral responses (saccades) was equal across all trials in our task, the monkeys faced perceptual challenges such as detecting very subtle stimulus changes on some trials. In our results, we demonstrated that LC neurons activated equally for all difficulties of stimulus changes, whether subtle or obvious. In the context of previous findings, we can conclude that although LC neurons encode information about the physical effort required for a behavioral response, they do not encode the amount of perceptual effort necessary for noticing or appropriately responding to a sensory stimulus.

Based on our results, future studies should consider the possibility that any observed stimulus-aligned LC responses are not purely sensory in nature, but instead may be linked to motor preparation processes triggered by the presentation of an important sensory stimulus, as we found in our study. Interestingly, all previous experimental paradigms used for studying LC phasic activation involved some variant of sensory cuing that could in principle trigger small eye, face, or body movements as preparations for impending, larger motor responses. For instance, in early studies of LC activity it is likely that startling or noxious stimuli prompted reflexive responses (e.g., muscle contractions, eye blinks associated with the startle reflex) or fidgeting movements not measured by experimenters. A previous study in rodents showed that LC phasic responses were associated with lipping (a Pavlovian reward expectation response) that occurred at the time of a reward cue and at the time of a task-related motor response (Bouret and Richmond 2009). However, few other studies of LC phasic activation have considered such potentially task-related small movements in their interpretations of sensory aligned responses. In future studies, it could be particularly illuminating to align LC activity on occurrences of small bodies (e.g., licking, fidgeting) and eye movements during task engagement, as some recent studies have done for cortical activity in rodents (Stringer et al. 2019). From the findings presented in this study, we conclude that the relatively brief, discrete saccade-aligned LC activations within seconds-long trials of our perceptual decision making task serve to orient and prepare the animal for an important behavioral response but have no impact on the perceptual accuracy of the animal’s decisions.

## Acknowledgements

This work was supported by NIH R01 MH128393. We are grateful to Sidd Joshi for helpful advice regarding the approach to locate and identify locus coeruleus neurons, to Richard Johnston for collaboration in behavioral training and mapping neuronal responses within the chambers, to Samantha Nelson for her technical and experimental support, and to our animal care staff.

---

**Figure 3 Supplement 1.**
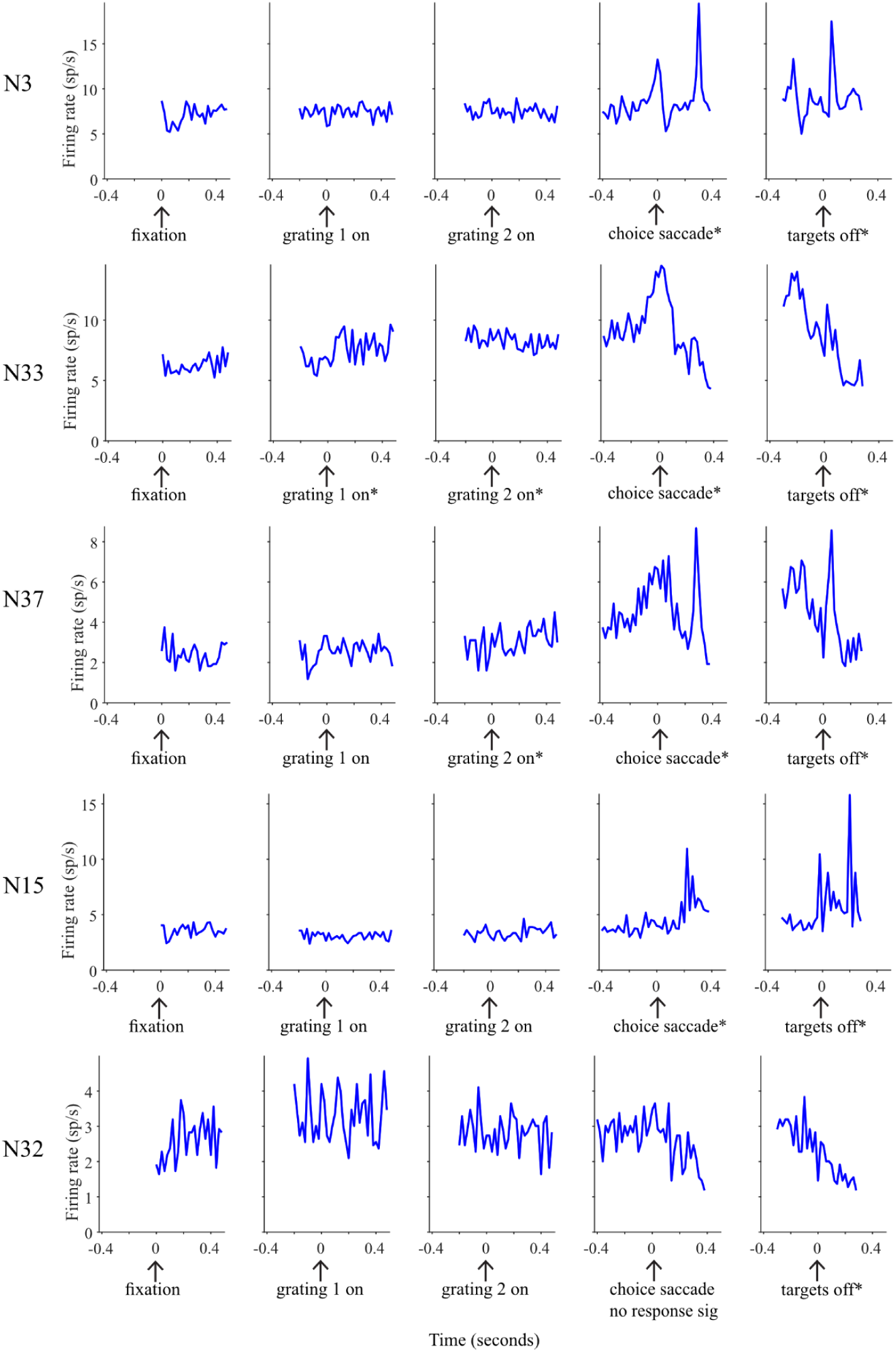
Example PETHs are shown for 5 neurons (rows) aligned to fixation (column 1), first stimulus grating onset (column 2), second stimulus grating onset (column 3), choice saccade initiation (column 4), and target offset (column 5). Neurons that had a significant change in response relative to baseline have an asterisk next to the plot label text for that condition.

**Figure 3 Supplement 2.**
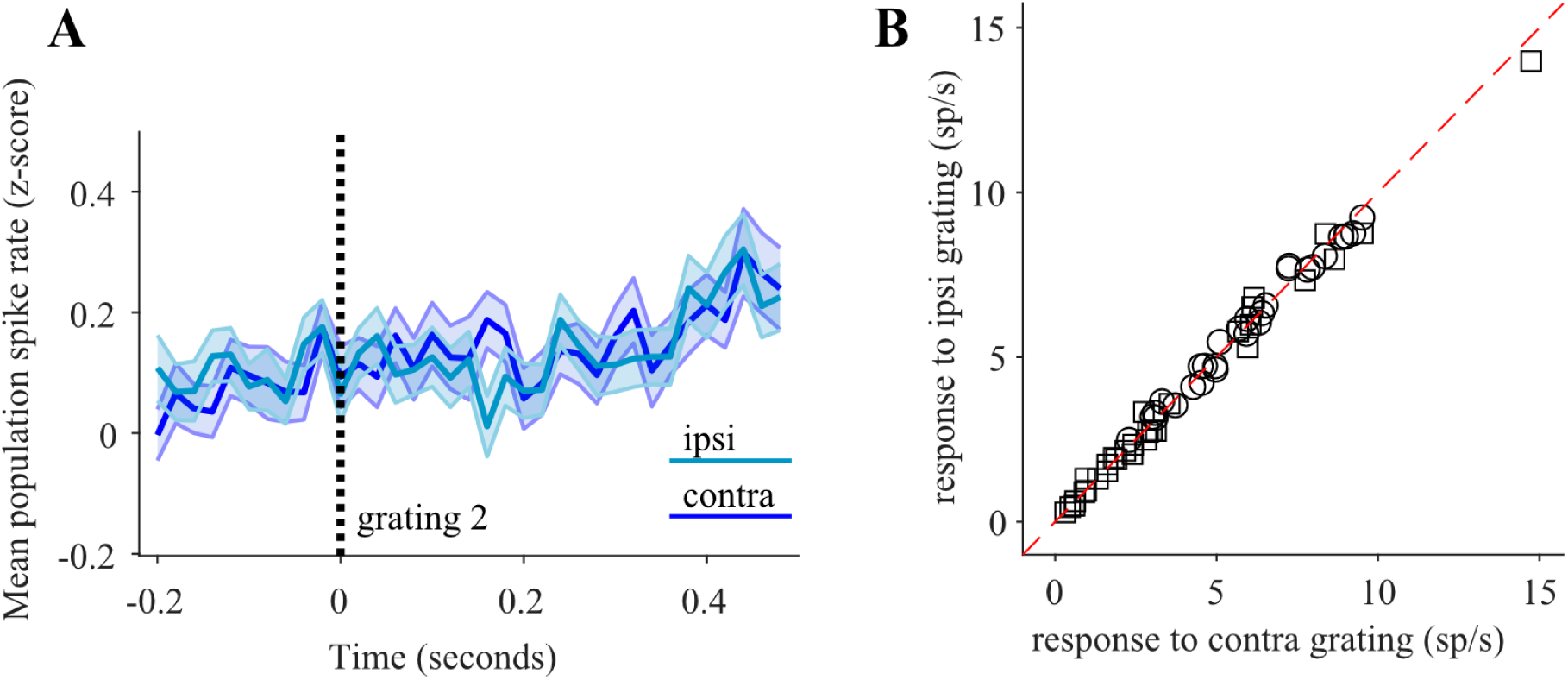
Task-related LC phasic activation during the test grating interval. (A) We did not observe any transient visual response to the appearance of the second grating, and (B) spike rates during the second grating interval showed no evidence of spatial selectivity based on whether the grating was presented ipsilateral or contralateral to the recorded LC neuron.

**Figure 4 Supplement 1.**
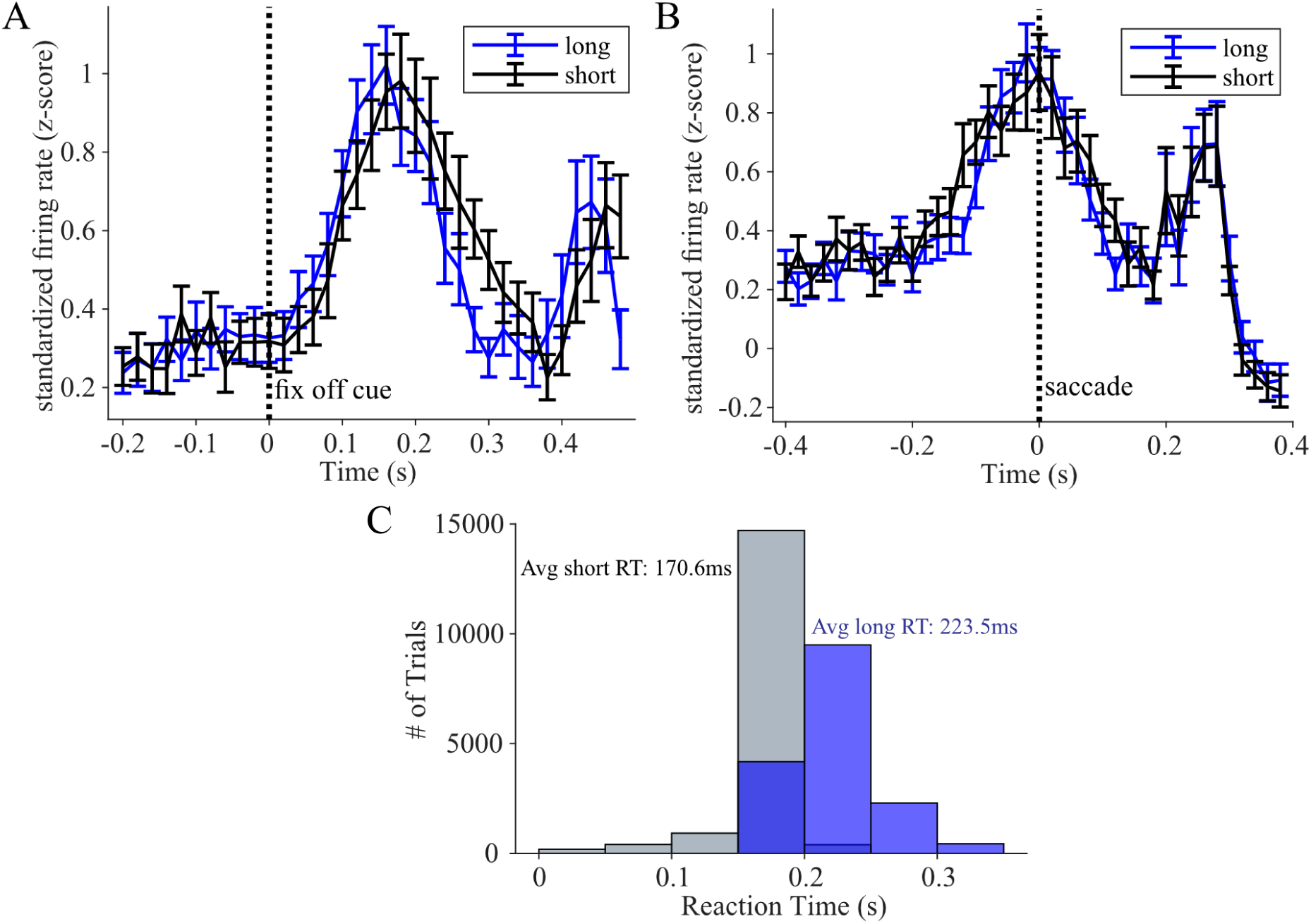
Comparing LC phasic activation between long and short behavioral response times. (A) Average PETHs compiled across choice target-onset aligned LC responses on long (blue) vs short (black) reaction time (RT) trials (lines and error bars represent mean ± SEM across PETHs of all individual neurons across both monkeys). Target-onset aligned LC firing rates were not significantly different between trials of short vs long RTs (paired t-test, p = 0.89). (B) same as (A) but for saccade-aligned LC responses; no significant difference between long and short RT trials (paired t-test, p = 0.5).(C) Histograms depict averages and distributions of behavioral response times in the short (gray) and long (blue) RT groups across sessions and monkeys. Long and short RT groups were determined by the median RT in each session.

**Figure 8 Supplement 1.**
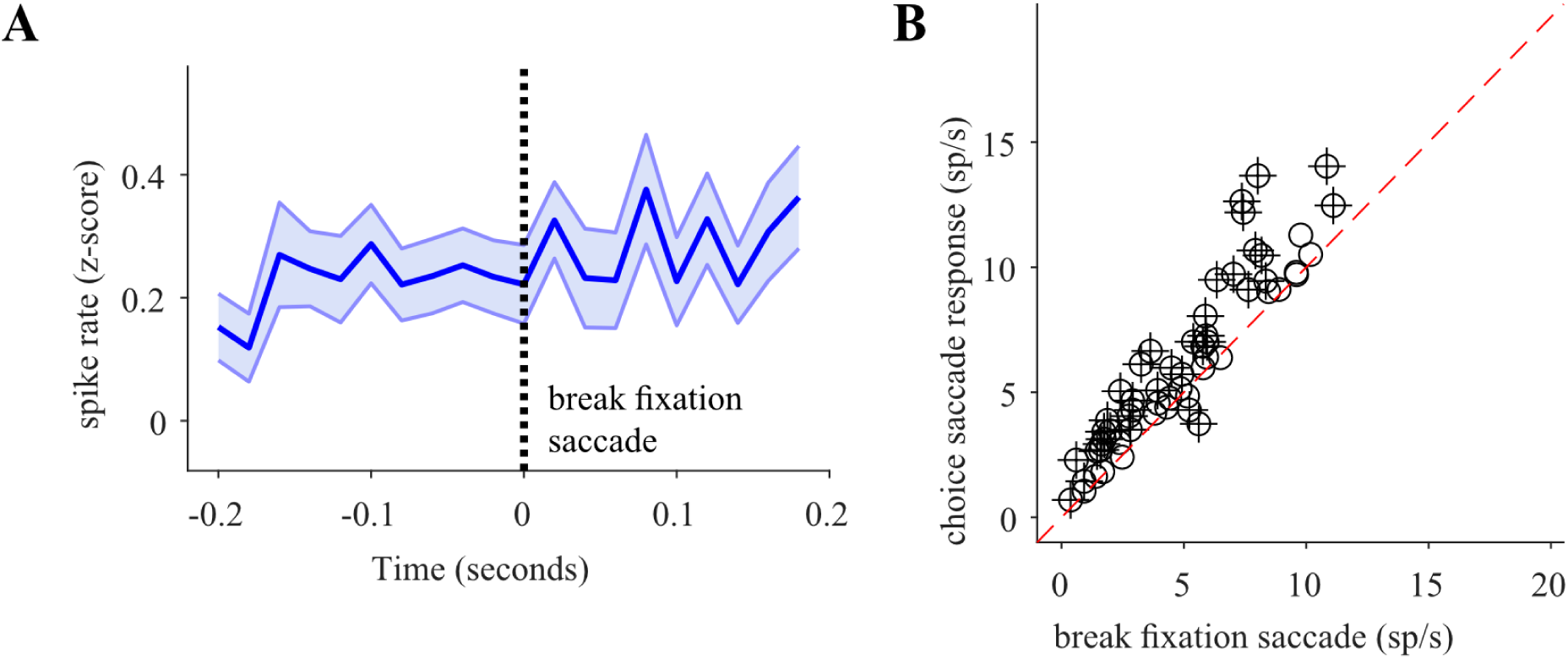
Relationship of LC activity to non-choice saccades. In addition to task-relevant behavior, in some cases the animals made a saccade away from fixation either to one of the gratings or elsewhere on the screen. These saccades were deemed task-irrelevant, were not rewarded and the trials were halted. (A) Average PETH across all standardized LC responses aligned on onset of task-irrelevant saccades that broke the animal’s fixation. (B) Scatter plot depicting the firing rate responses of 59 individual LC neurons aligned on the onset of the task-irrelevant break fixation saccades as compared to task-related choice saccades made during the choice period (y-axis). The majority of recorded neurons showed a significantly greater activation in response to task-related saccades (+ symbols indicate significant differences between the two responses, paired t-test, p < 0.05).

